# The Apurinic/Apyrimidinic Endodeoxyribonuclease 1 is an RNA G-quadruplex binding protein and regulates miR-92b expression in cancer cells

**DOI:** 10.1101/2024.02.22.581538

**Authors:** Alessia Bellina, Matilde Clarissa Malfatti, Gilmar Salgado, Aaron M. Fleming, Giulia Antoniali, Nicolò Gualandi, Sara La Manna, Daniela Marasco, Erik Dassi, Cynthia J. Burrows, Gianluca Tell

## Abstract

In the last decade, several novel functions of the mammalian Apurinic/Apyrimidinic Endodeoxyribonuclease 1 (APE1) have been discovered, going far beyond its canonical function as a DNA repair enzyme, unveiling its potential roles in cancer development. Indeed, it was shown to be involved in DNA G-quadruplex biology and RNA metabolism, most importantly in the miRNA maturation pathway and the decay of oxidized- or abasic-miRNAs during oxidative stress conditions. Furthermore, in recent years several non-canonical pathways of miRNA biogenesis have been described, with a specific focus on guanosine-rich precursors that can form RNA G-quadruplex (rG4) structures. In this study, we show that several miRNA precursors, dysregulated upon APE1-depletion, contain an rG4 motif and that their corresponding target genes are upregulated after APE1-depletion. We also show, both by *in vitro* assays and by using a HeLa cell model, that APE1 can bind and regulate the folding of an rG4 structure contained in pre-miR92b, with a mechanism strictly dependent on critical lysine residues present in the N-terminal disordered region. Furthermore, APE1 depletion in HeLa cells alters the maturation process of miR-92b, mainly affecting the shuttling between the nucleus and cytosol. Lastly, bioinformatic analysis of APE1-regulated rG4-containing miRNAs supports the relevance of our findings for cancer biology. Specifically, these miRNAs exhibit high prognostic significance in lung, cervical, and liver cancer, as suggested by their involvement in several cancer-related pathways.

**Significance Statement:** We highlight an undescribed non-canonical role of the mammalian Apurinic/Apyrimidinic Endodeoxyribonuclease 1 (APE1) in the context of RNA G-quadruplexes (rG4), specifically in the alternative pathway of miRNA maturation of guanosine-rich miRNA precursors. Specifically, APE1 binds these structures and modulates their folding, mainly through its N-terminal region and some residues in its catalytic domain. Moreover, we showed an interesting new role of APE1 in regulating the shuttling and accumulation of miR-92b between the nuclear and cytosolic compartments, opening new perspectives on how APE1 may exercise its role in the miRNA maturation pathway and function. Moreover, APE1-depleted dysregulated miRNAs with rG4 motifs in their precursors have significant prognostic value in lung, cervical, and liver tumors, suggesting potential targets for cancer therapy.

## Introduction

In humans, a large quota of RNAs is not translated into proteins and constitutes the non-coding fraction of RNAs (1). Within this class, microRNAs (miRNAs) and their precursors emerge for their increasingly recognized roles, being involved in several physiological and pathological processes (2, 3), as they mediate post-transcriptional gene expression suppression through the interaction with the Argonaute protein family (4). For their importance in cell physiology, miRNA maturation is finely regulated by enzymes and RNA-binding proteins (5). Their biogenesis starts with the transcription of a primary miRNA (pri-miRNA) by RNA polymerase II (6). The pri-miRNA folds into a stem-loop structure and is recognized and cropped by the Drosha-DGCR8 Microprocessor complex (5). Later, the resulting transcript, called precursor miRNA (pre-miRNA), is exported to the cytoplasm, where its terminal loop is cleaved by Dicer, releasing a small duplex RNA (7), which is then loaded onto the Argonaute protein and, after the removal of one of the two strands, becomes part of the mature RNA-induced silencing complex (RISC) to specifically target mRNA molecules based on the complementarity with the seed region (8, 9).

Recent studies have described an alternative pathway of miRNA maturation specifically tuned to guanosine-rich pre-miRNAs, which involves the existence of a thermodynamic equilibrium between the canonical stem-loop and a non-canonical secondary structure called RNA G-quadruplex (rG4) (10, 11). Analogously to DNA G-quadruplexes (G4s), rG4s are composed of stacks of guanine tetrads (called G-quartets) linked by Hoogsteen hydrogen bonds (12), mostly folded in a parallel topology with all strands following the same direction (13). Interestingly, this folded structure is more stable and compact than its DNA analogue, principally due to the 2’ hydroxyl group of the ribose (14). Several studies have shown that the presence of rG4s in miRNA may affect their maturation process (10, 11). Indeed, if present in pri-miRNAs, rG4s can inhibit Drosha-DGCR8 processing (15, 16); once present in pre-miRNAs, rG4s can inhibit Dicer activity (14, 17–21), whereas in mature miRNAs rG4s can impede loading onto the RISC complex (22–25). Therefore, the effect of this multi-level inhibition may lead to an aberrant miRNA maturation, impairing their ability to target mRNAs, thus potentially affecting several cellular pathways (26, 27). For these reasons, targeting rG4s in miRNAs and their precursors could be explored as a novel therapeutic strategy. A very well-characterized example is represented by pre-miR-92b, which is structured to form a stable rG4 at physiological KCl concentrations (28). Mature miR-92b is clinically relevant, being overexpressed in several human cancers including cervical and gastric cancers, glioma, and non-small cell lung cancer (NSCLC) (29–32). It was shown that the stabilization of its rG4 by a locked nucleic acid resulted in a decreased maturation of the respective miRNA, leading to the rescue of the expression of its target PTEN in NSCLC cells, thus sensitizing cancer cells to doxorubicin treatment (33). APE1/Ref1 (Apurinic/Apyrimidinic Endodeoxyribonuclease/Redox Effector Factor 1) is a multifunctional protein, mostly known for its role as endonuclease in the base excision repair (BER) pathway in the repair of oxidative DNA lesions and functioning as a redox hub for many transcription factors (TFs) (34). In recent years, great attention has been paid to the fascinating non-canonical roles of APE1 (34, 35). Indeed, APE1 mediates transcriptional regulation by stabilizing G4s at promoter regions, creating a platform for TF recruitment (36–38). *In vitro* experiments have demonstrated that APE1 maintains its endonuclease activity towards abasic sites embedded in G4s, although with a lower efficiency when compared to duplex DNA (39, 40). Surprisingly, the N-terminal region of APE1 is essential not only for G4s binding but also for modulating the catalysis of the abasic site cleavage. Remarkably, those activities largely depend on the ionic strength of the reaction, and above all on the magnesium concentration (37, 39). Furthermore, the N-terminal portion of the protein presents several lysine residues, most specifically 27, 31, 32, and 35, whose acetylation increased the cleavage ability of the protein and diminished the capacity to bind chromatin, due to the neutralization of its positive charges (39). Most interestingly, APE1 is involved in oxidized- or abasic-pri-miRNA processing and stability (41) during oxidative stress. Data from our laboratory have demonstrated that APE1 efficiently binds to pri-miR-221 and pri-miR-222 in HeLa, HCT-116, and MCF-7 cell lines and that their processing is affected in APE1-depleted cells, resulting in an upregulation of PTEN, a target gene of both miRNAs, during oxidative stress conditions. Moreover, we showed that APE1 interacts with Drosha but not with DGCR8 and DDX5, suggesting that it may affect the early phases of the microprocessor pathway and may compete with Dicer for the processing of some pre-miRNAs (42). Lastly, APE1 is involved in controlling Dicer expression in NSCLC through APE1-regulated miRNA-33a-5p and miRNA-130b (43). From several studies performed in the last thirty years by different groups, it is now clear that APE1, which is overexpressed in the nucleus and cytoplasm of many tumor types (e.g. hepatic, cervical and lung), is involved in tumor progression (44–48), thus representing an important target for cancer therapy (35). However, the molecular mechanisms responsible for the tumorigenic role of APE1 are still unclear but point to the non-repair activities of the protein, such as redox regulation and RNA metabolism. Our recent data suggest that APE1 may contribute to the expression of chemoresistance and tumor progression genes through post-transcriptional mechanisms involving miRNA regulation (49–51, 41). However, whether APE1 is endowed with the ability to recognize oncogenic miRNAs in a sequence-specific manner, or whether rG4 structures may play a role in its binding properties are completely unknown and could contribute to understanding the role of APE1 in the tumorigenic process.

Therefore, this study aims to investigate the involvement of APE1 in the stabilization/destabilization effect of pre-miRNAs containing rG4 motifs, as this protein might regulate the equilibrium between the G-quadruplex and the canonical stem-loop structures, thus impacting miRNA processing.

Through bioinformatic analysis, performed on a list of differentially expressed miRNAs obtained from HeLa and A549 cells upon stable downregulation/reconstitution of APE1 protein, we investigated the presence of rG4s in pre-miRNAs, finding that about 27% of those precursors contain an rG4-forming motif. *In vitro* experiments, using a synthetic oligoribonucleotide mimicking the rG4-containing pre-miR-92b, demonstrated that APE1 can bind and affect the folding and stability of this structure. In addition, we analyzed the region of APE1 that binds rG4 motif contained in pre-miR-92b and compared this region with previously published contact surfaces between APE1 and different enzymatic targets such as DNA. Furthermore, we confirmed these data in HeLa cells, showing that APE1 can bind pri-miR-92b, and that this is mainly mediated by the lysine residues present in its unstructured N-terminal region. Finally, we observed that miR-92b subcellular localization between nuclei and cytosol is deeply influenced by APE1 expression, thus suggesting that APE1 could be an important regulator of miR-92b maturation and shuttling processes. A bioinformatic analysis of the predicted rG4-containing miRNAs regulated by APE1 highlighted their prognostic significance in lung, cervical, and liver tumor types. At the end, by functional enrichment analysis, we reported the involvement of these miRNAs in several cancer-related pathways, such as epithelial to mesenchymal transition, tumor progression, and immune-evasion. We thus provide evidences for a novel layer of APE1 functions, which could mediate its role in cancer development.

## Results

### APE1 depletion causes a dysregulation of rG4 motif-containing pre-miRNAs

Starting from a list of miRNAs dysregulated upon APE1 depletion (n=248, Dataset S1) obtained by Nanostring and RNA-seq analysis in HeLa and A549 cell lines (41, 43),(Mangiapane G. et al., 2023, under revision), we searched for their corresponding pre-miRNAs (n=227) by using miRbase (52). Then, we examined the presence of rG4-forming motif sequences in the pre-miRNAs through a bioinformatic analysis using three algorithms, namely QGRS mapper (score threshold = 19, (53)), pqsfinder (score threshold = 47, (54, 55)) and G4Hunter (score threshold = 1,2, (56, 57)). In total, 61 of the initial 227 sequences were predicted to contain an rG4 motif at least by one tool, which indicated that about 27% of the pre-miRNAs, dysregulated by APE1 depletion, potentially contained an rG4 motif (Fig. 1A, Dataset S2). Among 61 pre-miRNAs, 15 sequences were predicted by all three algorithms (6.6%) with above-threshold scores; 19 were predicted by both QGRS mapper and psqfinder (8.4%), while the remaining 27 were predicted by only one of the three tools (12%). All these data confirmed the similar prediction obtained by Arachchilage and colleagues where about 16% of human pre-miRNAs hold a rG4 motif (28). On our side, we further observed that following APE1 depletion, the number of dysregulated precursors of miRNA holding rG4-forming sequences was higher, leading us to the hypothesis that APE1 has a molecular role in the regulation of rG4-containing miRNAs (Fig. S1A and Fig. 1B).

**Figure 1:**
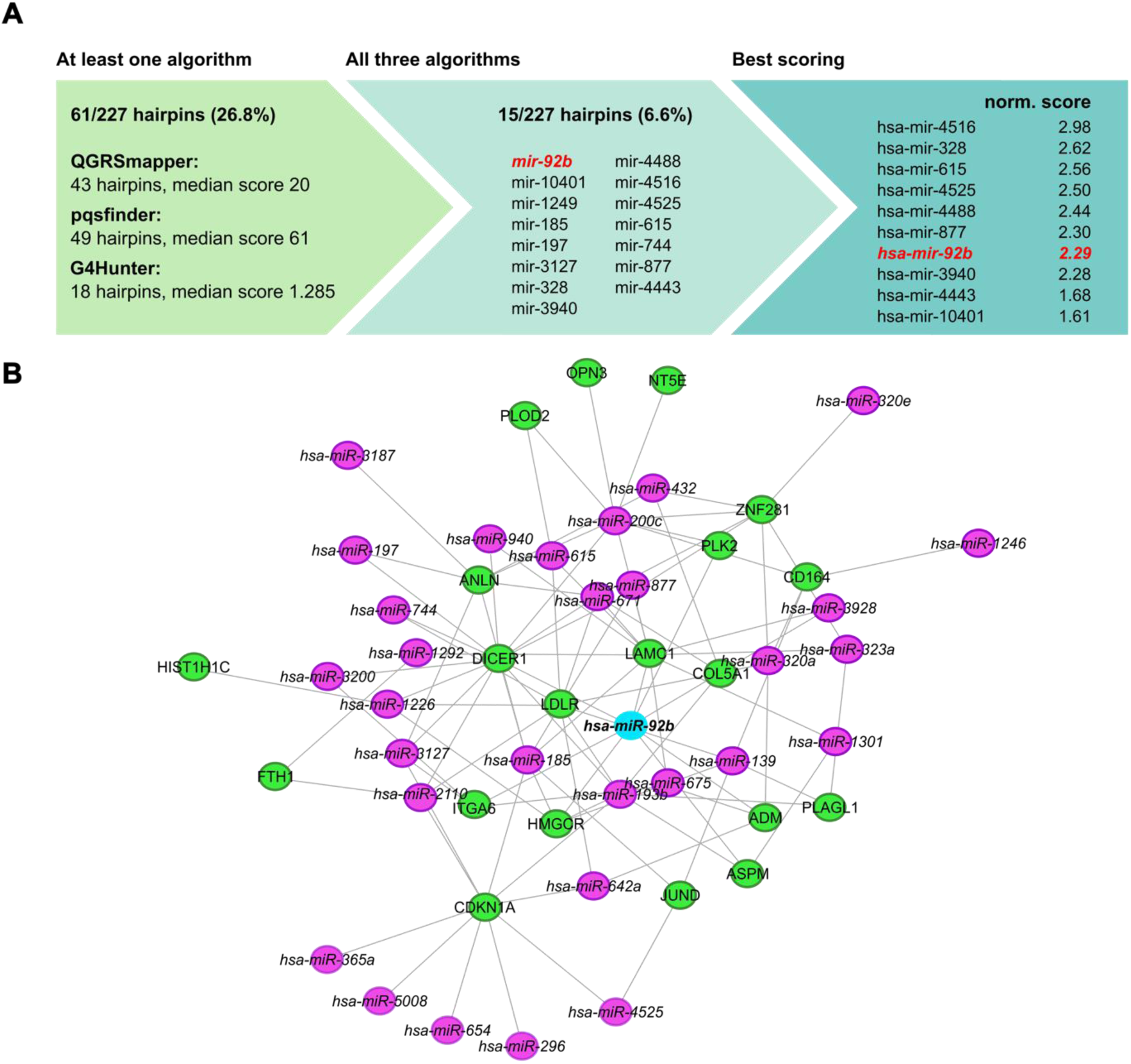
Predicted rG4-containing pre-miRNAs dysregulated upon APE1 silencing control several upregulated genes associated with APE1 depletion. A) Number of pre-miRNAs with at least a predicted rG4 for each algorithm and the related median prediction score (left); the list of pre-miRNAs predicted to have at least an rG4 by all three algorithms (center); and the top ten pre-miRNA ranked by their normalized prediction score, computed as the sum of each algorithm’s score scaled by the maximum obtained score for that algorithm (right). B) Network of miRNA-gene interaction for the 61 miRNA present in the signature. Only upregulated genes (p-value < 0.05 and log2FoldChange > 0) in both of the cell lines analyzed (HeLa or A549) were reported. Each purple node represents a miRNA while the edges represent a miRNA-gene interaction reported in Tarbase. Upregulated genes are reported as green nodes.

Next, to identify the pathways associated with genes upregulated following APE1 depletion, we performed a pathway enrichment analysis using the KEGG database (58). Genes exhibiting upregulation were defined as those with an adjusted p-value < 0.05 and a positive log2(Fold Change) in APE1-depleted cells, compared to control cells, in at least one of the two cell lines examined (HeLa or A549). Our findings revealed that the upregulated genes primarily participate in oncogenic pathways such as the “p53 signaling pathway”, “Pancreatic cancer”, “Transcriptional misregulation in cancer”, “Small cell lung cancer”, “Proteoglycans in cancer” and “Colorectal cancer”, further confirming the role of APE1 in cancer onset and progression (59). Notably, several upregulated genes were significantly enriched in the “Focal adhesion” pathway, implicating a role of APE1 in processes related to extracellular matrix organization and cell-to-cell interactions, whose dysregulation could lead to different human diseases including cancer. Indeed, focal adhesion proteins contribute to multiple aspects of cancer, including: increased cell proliferation, resistance to apoptosis, elevated cell motility and invasion, and promotion of angiogenesis (60). In line with this result, we also identified significantly enriched pathways related to senescence and the cell cycle, reinforcing the role of APE1 in those processes (Fig. S1A). Finally, to explore the relationship among upregulated genes and dysregulated rG4-containing miRNAs upon APE1-depletion, we built a miRNA-gene interaction network (Fig. 1B) reporting the Tarbase-validated interactions between each of the 61 rG4-containing miRNA and the commonly upregulated genes in both HeLa and A549 cells lines after APE1-depletion (61). Notably, we observed the presence of different connections between rG4-containing miRNAs and upregulated genes following APE1-depletion. Intriguingly, miR-92b, located in the center of the network, directly and indirectly regulates the expression of many upregulated genes suggesting its pivotal role in the regulation of the expression of key genes associated with APE1-depletion (41) (Fig. 1B).

### APE1 binds rG4-pre-miR-92b and induces conformational changes

G4 motifs are known to be present in miRNAs and their precursors (10). Given the role of APE1 in miRNA processing and in the regulation of G4 structures (37, 39–41, 43), we investigated whether APE1 might bind and modulate the folding of rG4 structures in miRNAs.

To assess this hypothesis, we chose pre-miR-92b as a general model for our next investigations for three main reasons: (i) pre-miR-92b was predicted to contain an rG4 by all three tools with high scores and controls target genes already known to be regulated by APE1 (i.e. PTEN, DICER; CDKN1A); (ii) pre-miR-92b contains an rG4 motif already experimentally validated and characterized (28), and (iii) the stabilization of its rG4 represents an attractive therapeutic target (28, 33). Based on the QGRSmapper analysis (reported in Table S1), we first designed an rG4-forming sequence derived from pri-miR-92b and pre-miR-92b sequences (Fig. 2A). This sequence, hereafter named pre-92b and highlighted in green, is entirely located in pre-miR-92b, close to the Drosha cleavage site (Fig. 2A). This sequence was used for the following *in vitro* analyses with or without a fluorophore in the 5’-end. In our first structure analysis, the non-fluorescent oligoribonucleotide was folded in a 50 mM K^+^ solution and then inspected by CD (Fig. 2B) and NMR (Fig. S1B) spectroscopies. The CD spectrum showed a positive λ_max_ = 263 nm and a negative λ_min_ = 242 nm molar ellipticities, consistent with a parallel-stranded rG4 (Fig. 2B) (62), as confirmed by 1-D NMR spectrum, indicating the formation of several imino peaks compatible with two or more conformers of three G-tetrad rG4 in dynamic equilibrium (Fig. S1B).

**Figure 2:**
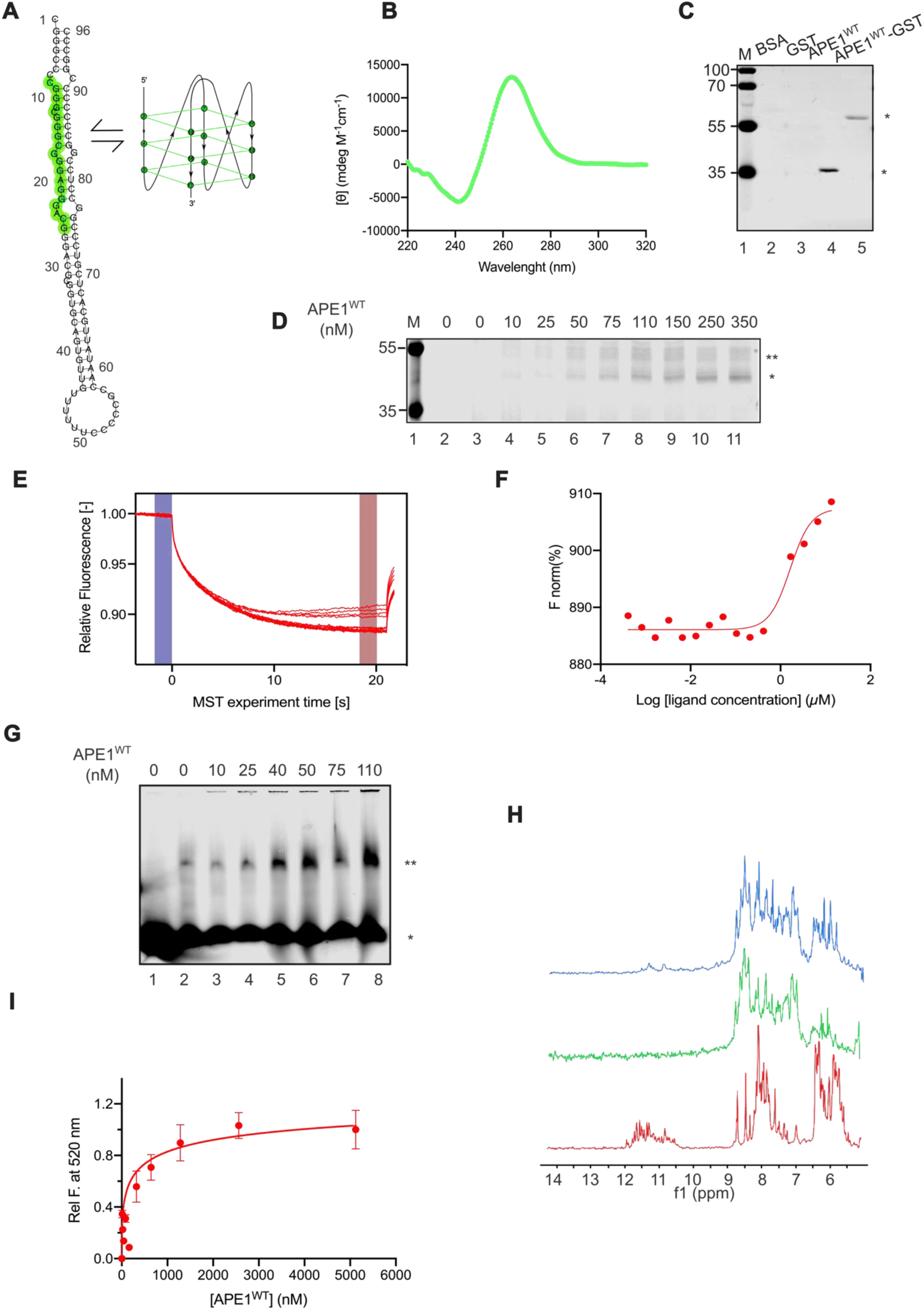
APE1 binds pre-miR-92b rG4 and regulates its folding equilibrium. A) pre-miR-92b structure in which the rG4 sequence studied is highlighted in green (pre-92b). On the right, the graphic scheme of the parallel rG4 is formed by the highlighted sequence. B) CD spectrum for the pre-92b probe demonstrates its folding into a parallel-stranded rG4. The wavelength (expressed in nm) and the CD (expressed in mdeg M^-1^ cm^-1^) are indicated on the x- and y-axis, respectively. C) Representative NWB shows the binding between pre-92b probe and recombinant APE1^WT^ untagged and tagged with GST (APE1^WT^-GST). BSA and GST were used as controls. 1500 ng of each protein was loaded on the SDS-PAGE gel,blotted on the membrane and then incubated with the pre-92b fluorescent probe (5 pmol). On the left, the electrophoretic marker is loaded, and the different molecular weights are expressed in kDa. D) Representative crosslinking analysis with pre-92b probe (25 nM) and different amounts of APE1^WT^ reported upon the image and expressed in nM. On the left, the electrophoretic marker is loaded and the different molecular weights are expressed in kDa. In lane 2, the oligoribonucleotide was heated at 70°C before loading, as control. E) Thermophoretic traces of the MST assay between pre-92b and APE1^WT^, with the relative fluorescence (expressed in [-]) on the y-axis and the MTS experiment time on the x-axis (expressed in s). F) Binding isotherm of the MST signals *versus* the logarithm of APE1^WT^ concentrations (µM). G) REMSA analysis with different amounts of recombinant APE1^WT^, reported upon the gel and expressed in nM, incubated with pre-92b probe (25 nM) in 100 mM KCl. The asterisks (*) and (**) indicate the unbound probe bands and the oligomeric band, respectively. In lane 1, the protein-free oligoribonucleotide was heated at 70°C before loading, as control. H) Proton NMR spectra depicting 100 µM pre-92b in 10 mM KPi buffer with KCl and 0.5 mM MgCl_2_. In red, the heterogenous folding pattern of different rG4 in a complex equilibrium. Green depicts the pre-92b after incubation with 1 molar equivalent of APE1. Blue spectrum shows the 1:1.2 molar ratio of protein to pre-92b. I) Fluorescence study to monitor the structural conversion of an RNA duplex to an rG4 structure facilitated by recombinant APE1 protein (expressed in nM). The APE1^WT^ concentration (expressed in nM) and the Relative Fluorescence at 530 nm are indicated on the x- and y axis respectively.

With the rG4 folding of the pre-miR-92b confirmed in these experimental conditions, we evaluated the ability of APE1 to bind the rG4 motif. First, we carried out a NWB assay by analyzing equal amounts of BSA, GST, recombinant purified APE1 wild-type (APE1^WT^), as well as the tagged APE1^WT^-GST protein (Fig. S1C). After gel running and blotting, we renatured the proteins by incubation with serial dilutions of GdnHCl and then we incubated with the soluble fluorescent probe. As shown in Fig. 2C, we observed that both APE1^WT^ and APE1^WT^-GST (lanes 4 and 5) were able to bind the probe, as indicated by the asterisk. The absence of any bands in the BSA and GST lanes (lanes 2 and 3) confirmed that the binding of the probe to APE1^WT^ was highly specific.

To confirm these data, we carried out a UV-crosslinking assay using increasing amounts of APE1^WT^ protein and a constant amount of pre-92b probe (Fig. 2D). As indicated by the single asterisk, the observed band corresponded to the UV-crosslinked complex between APE1 and the probe. Specifically, APE1 was able to bind the probe in a dose-dependent manner (lanes 4-11), forming a 1:1 ratio complex of 43 kDa. Some higher molecular complexes were also observed as faint bands and highlighted by double asterisks. As expected, all these bands were not observed in the protein-free oligoribonucleotide kept at room temperature (lane 3) and in the protein-free oligoribonucleotide heated at 70°C before loading (lane 2).

To quantitatively evaluate the affinity between pre-92b probe and APE1^WT^, we performed a MST assay (Fig. 2E and 2F). Keeping the pre-92b probe concentration constant and increasing the APE1^WT^ concentration, we obtained a dose-response profile and the fitting of the data allowed us to estimate an IC_50_ value of 1.6 ± 0.6 μM,.

Since was already known that APE1 facilitates the conversion of DNA-duplex structures to G4s (37), to characterize the possible involvement of APE1 in the folding process of rG4s, we used REMSA analyses (63) with the recombinant purified protein. As shown in Fig. 2G, in the presence of KCl, a retarded band at low intensity was observable in the protein-free probe reaction (lane 2), suggesting the possible presence of slow-migrating intermolecular secondary structures, maybe due to oligoribonucleotide dimer/oligomer formation. Indeed, this shifted band was not present under denaturing conditions (by heating the sample at 70°C) (lane 1). Interestingly, when we incubated the probe with increasing concentrations of APE1^WT^, the intensity of the retarded band increased as a function of APE1 concentration, suggesting that APE1 could have a potential role in changing the equilibrium between monomer and dimers/oligomers (lanes 3-8, indicated by **), as we already observed with the telomeric G-rich sequence (39). A similar experiment performed with a polyU oligonucleotide of the same length confirmed the specificity of APE1 for the rG4 containing sequence (data not shown). The same assay was carried out in the LiCl buffer (Fig. S1D), which is known to destabilize G4 formation (64). Indeed, the low-intensity band in the protein-free lane was far less apparent in the presence of LiCl (lane 5) compared to KCl buffer (lane 2). APE1 seemed to improve the intensity of the retarded band in a dose-dependent manner, both in the presence of KCl (lanes 3 and 4) or LiCl (lanes 6 and 10), bypassing the destabilizing effect of LiCl.

We then inspected the hypothesis for a role of APE1 in modulating the folding equilibrium of the rG4 structure with a dedicated structural technique by analyzing the structure of the complex between APE1 and the oligoribonucleotide by solution-state NMR spectroscopy.

The ^1^H NMR spectrum (Fig. 2H) of 100 µM of pre-92b oligoribonucleotide in KPi buffer was recorded at 25°C. The spectrum highlighted in red evidenced a complex imino peak pattern typical of G4s. We could observe numerous imino peaks in the region of 10-12.5 ppm, indicating a heterogeneous folding pattern of different G4s in a complex equilibrium. The green spectrum depicted the pre-92b probe after incubation with 1 molar equivalent of APE1 for one hour before recording the same number of transients. The imino fingerprint vanished in the presence of APE1, indicating either an unfolding of the rG4 into a single-stranded molecule, which was in fast exchange with the water solvent and did not allow the formation of stable imino bonds between the nucleobases, or potentially the formation of higher order aggregates could explain this phenomenon. When we added further oligoribonucleotide (1:1.2 total molar ratio) to the NMR tube (blue spectrum), we could observe the reappearance of some imino peaks, indicating a saturation level and a stable complex of 1:1 interaction of APE1 with G-rich RNA in which APE1 modulates the folding behavior of the rG4.

Finally, to understand whether APE1^WT^ could affect the folding of the possible rG4 sequence in pre-92b from a duplex-like RNA structure, we performed a fluorescence spectroscopy analysis. To conduct this experiment, a 3′-FAM-labeled pre-92b probe was annealed with a complement bearing a 5′ quencher (BHQ), and the sequence provided the two natural bulges predicted in the pre-mir-92b structure (Fig. S1E). The protein was titrated into the RNA solution while monitoring the evolution of the FAM fluorescence as the structure switched from duplex to rG4 (Fig. 2I). The experiment identified that APE1^WT^ could facilitate refolding of the duplex to an rG4 with a midpoint of 400 ± 90 nM (Fig. 2I, red line), as happens for G4-DNA (37).

In summary, our data confirm the hypothesis that APE1 can bind rG4-containing pre-miR-92b and modulate the folding equilibrium of its rG4 structure.

### Lysine residues within the APE1 N-terminal region are required for stable binding to pre-miR-92b

Recent works from our laboratory and others showed that APE1 binding to G4-DNA, both in promoters and telomeres, highly depends on its N-terminal region (37, 39). To deepen the role of this region in the recognition mechanism of the rG4 sequence, we performed an *in vitro* assay using recombinant purified APE1^WT^, together with APE1^NΔ33^ mutant protein, lacking the first 33 N-terminal residues, and APE1^K4pleA^ protein, a mutant in which lysine residues 27, 31, 32 and 35 are replaced by alanine residues to reset the positive charges of the lysine side chains (65). First, we performed a NWB assay (Fig. 3A) by loading equal amounts of APE1^WT^, APE1^NΔ33^, and APE1^K4pleA^ recombinant proteins and BSA, as a negative control (Fig. S2A). As expected, APE1^WT^, APE1^NΔ33^ and APE1^K4pleA^ were able to bind to pre-92b probe with different strengths (Fig. 3A); indeed, the APE1^NΔ33^ quantified band was weaker than those of APE1^WT^ and APE1^K4pleA^. Specifically, APE1^NΔ33^ lost half of its binding ability toward the pre-92b probe compared to APE1^WT^, while APE1^K4pleA^ showed a 30% reduction of its binding ability (Fig. 3B).

**Figure 3:**
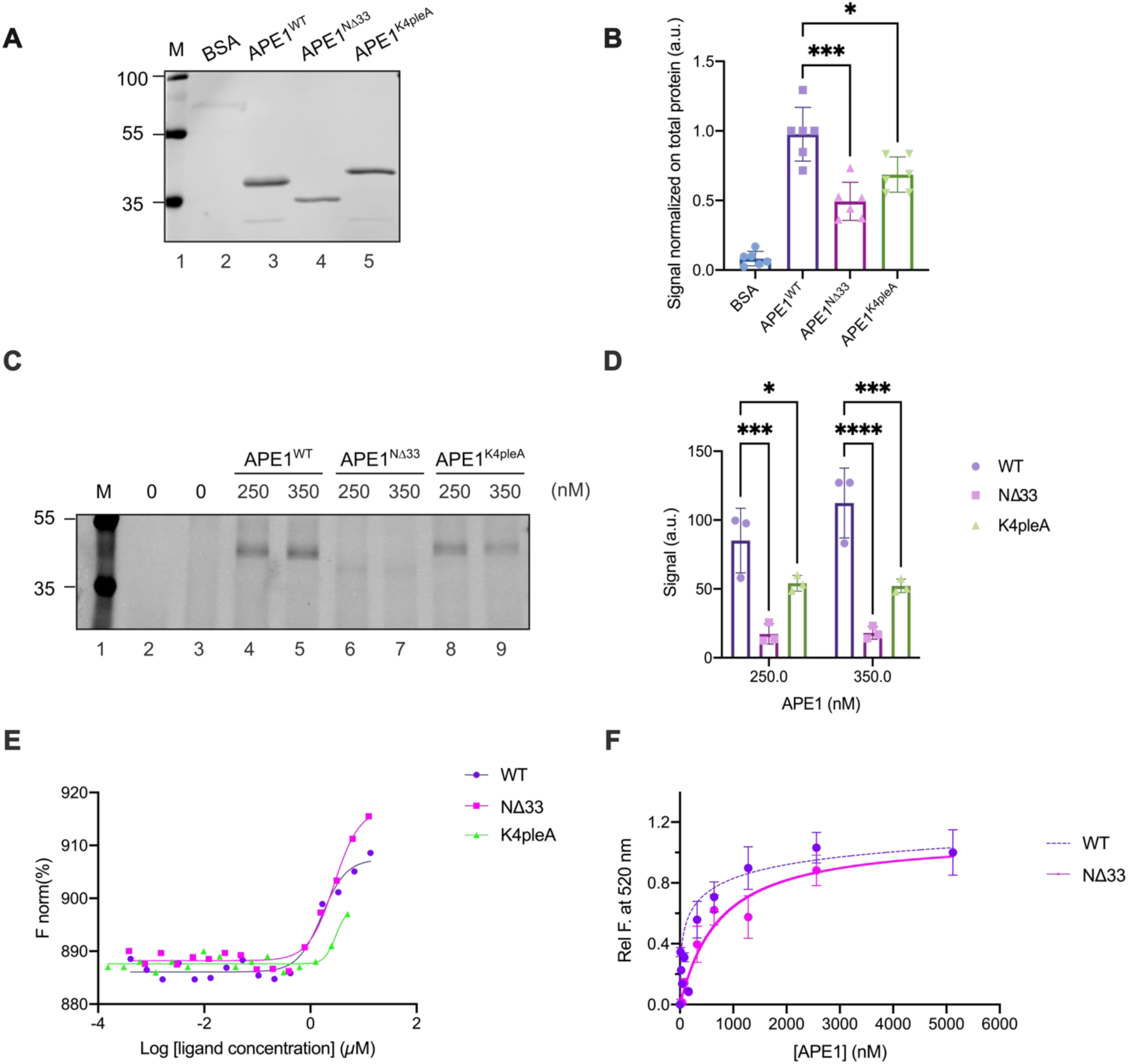
APE1 N-terminal region is necessary for pre-miR-92b rG4 binding. A) Representative NWB shows the binding between pre-92b and recombinant APE1^WT^, recombinant APE1^NΔ33^ and recombinant APE1^K4pleA^. BSA was used as a control. 1 µg of each protein was loaded and blotted on the membrane and then incubated with the pre-92b (5 pmol). On the left, the electrophoretic marker is loaded, and the different molecular weights are indicated in kDa (MW). B) Relative histogram summarizing the different binding abilities towards pre-92b between BSA, APE1^WT^, APE1^NΔ33^, and APE1^K4pleA^, normalized on the respective total protein staining signal (Fig. S1 B). The data are expressed as means ± S.D. of six independent replicates. C) Representative crosslinking analysis of with different amount, expressed in nM, of APE1^WT^, APE1^NΔ33^, APE1^K4pleA,^ recombinant proteins and pre-92b probe (25 nM), On the left, the electrophoretic marker is loaded, and the different molecular weights are expressed in kDa. D) Relative histogram summarizing the different binding abilities towards pre-92b between APE1^WT^, APE1^NΔ33,^ and APE1^K4pleA^, at two different doses (250 and 350 nM). The data are expressed as means ± S.D. of three independent replicates. E) Overlay of the isotherms of MST assays between pre-92b and the logarithm of APE1^WT^, APE1^NΔ33^ and APE1^K4pleA^ concentrations (µM). The binding isotherm for the MST signal between pre-92b *versus* APE1^WT^, APE1^NΔ33,^ and APE1^K4pleA^ concentrations is reported. F) Fluorescence study to monitor the structural conversion of an RNA duplex to an rG4 structure facilitated by recombinant APE1 and APE1^NΔ33^ proteins (expressed in nM). The proteins concentration (expressed in nM) and the Relative Fluorescence at 530 nm are indicated on the x- and y axis respectively. A p-value < 0.05 is symbolized by a single asterisk (*), while a p-value < 0.001 is symbolized by three asterisks (***) and < 0.0001 by four asterisks (****).

Subsequently, as previously done for APE1^WT^, we investigated if APE1^NΔ33^ and APE1^K4pleA^ could form a stable rG4-protein complex. A UV-crosslinking assay was carried out using APE1^NΔ33^ and APE1^K4pleA^ in comparison with APE1^WT^ (Fig. 3C). Differently from what we observed for APE1^WT^, the reactions between the probe and APE1^NΔ33^ and APE1^K4pleA^ provided very weak bands, both corresponding to the crosslinked adducts.The different rG4-binding abilities of the three proteins at the two different doses considered were quantified and plotted in the graph shown in Fig. 3D. Interestingly, APE1^NΔ33^ lost almost 80% of its rG4-binding ability, while the reduction observed in the case of APE1^K4pleA^ mutant was close to 50%.

To confirm these data, we conducted a MST assays by using all above mentioned APE1 proteins (Fig. 3E, Fig. S2B and Fig. S2C). The fitting of the dose-dependent signals provided an IC_50_ value of 2.8 ± 0.3 μM in the case of APE1^NΔ33^ mutant protein. Conversely, in the case of APE1^K4pleA^, the fitting did not converge due to the lack of saturation values since this construct exhibited a limited water solubility with respect to the others. To deepen potential differences in the biophysical behaviors of APE1 constructs, we evaluated their CD spectra upon temperature increase. While APE1^WT^ and APE1^NΔ33^ exhibited a cooperative unfolding process providing similar T_m_ values (∼ 44°C, (66)) (Fig. S2D, Fig. S2E, and Fig. S2F), in the case of APE1^K4pleA^ we did not observe a cooperative mechanism, but rather the occurrence of aggregation upon temperature increase (Fig. S2F). This aggregation propensity could explain the lower solubility. Finally, we used a fluorescence analysis to evaluate if the N-terminal truncated mutant could regulate the rG4 folding from a duplex-structure, as happens for APE1^WT^ (Fig. 3F, pink line). The experiments showed that APE1^NΔ33^ could facilitate refolding to the rG4 at a 2-fold higher concentration than APE1^WT^ (780 ± 110 nM vs. 400 ± 90 nM), demonstrating that the N-terminal region of the protein is necessary for the rG4 folding regulation of pre-92b probe.

Together, these data indicate that the N-terminal region of APE1 exerts a key role in the binding and regulation of rG4 structures.

### APE1 interacts with pre-92b sequence through its canonical DNA binding interface

To investigate the interaction between APE1 and the pre-92b RNA molecule, we performed 2D NMR spectroscopy. ^15^N-^1^H TROSY-HSQC experiments were performed with 60 µM of APE1 (Fig. S3A) alone (green) and in the presence of 60 µM of pre-92b probe (red). As already observed in previous published NMR and X-ray crystallography experiments for the apo-APE1 (67) and APE1-DNA complex structures (68), the N-terminal region of the protein is not resolved due to highly flexible and intrinsically disordered amino acids. Nevertheless, using the 2D NMR spectra, we were able to follow important chemical shift modifications to key residues during the titration with pre-92b probe. In the apo form (green spectra), most residues were correctly identified, with aid from BMRB entry code 16516. Upon the addition of one molar equivalent of the pre-92b probe, prominent modifications in the spectrum occurred. Those chemical shift modifications were categorized into 3 groups: the first group, composed of residues that disappeared after the addition of RNA, a second group that were not visible in the apo form while they appeared in the spectrum after RNA inclusion (red overlay spectra), and the third group, which corresponded to visible peaks in the apo form that were shifted during the titration due to chemical shift changes. The disappearance of the residues can be explained by the extensive line broadening during the titration and as the result of moderate affinity (µM range) between those amino acids and the RNA molecule that are usually in the range of intermediate exchange regime (k_ex_ ≈ |Δω|). Between the disappearing residues localized in the canonical DNA binding region of APE1, we could find E216, T265, F266, M271, S307, and H309. For the second group, we observed the appearance of peaks at stoichiometric amounts, which led us to infer that the RNA molecule could induce conformational exchange processes. Indeed, we observed peaks for residues localized in dynamic regions or undergoing conformational exchange, which were stabilized by interactions such as hydrogen bonds or salt bridges, often associated with specific binding. These peaks corresponded to amino acids directly involved in the ligand binding site, such as Y262 and M270. A few more peaks were observed but they could not be assigned unambiguously. We then plotted the disappearing and reappearing peaks, and the most significantly shifted peaks in the overlay of the apo (Fig. S3B, peaks depicted in red) and the bound form (Fig. S3C, peaks depicted in red) of APE1, with respective PDB accessing codes of 1BIX and 1DEW. The structural representation allowed us to clearly identify the involvement of the helix-hairpin-helix (HhH) motif, which is responsible for recognizing and binding many DNA targets, including G4s (36). Among the different amino acids that make part of the recognition and enzymatic site, apo-APE1 contains two important residues that were well-conserved and undergo structural changes upon nucleic cid binding, e.g., R177 and M270 (cyan). These residues form a bridge-like structure that can be inserted into the major and minor grooves of RNA, respectively (purple for R177 and yellow for M270).

Although we could not unambiguously identify R177 in the spectrum, M270 reappearance made us believe that APE1 binding towards the pre-92b probe is compatible with a classic binding mechanism observed in many other targets (69). For comparison, we left the DNA structure (blue ribbon) of the protein-DNA complex identified with PDB code “1DEW”.

### APE1 regulates the maturation and shuttling of miR-92b in cells with a high prognostic value in real-world cancers

Once we established that APE1 can bind and regulate rG4 folding in pre-miR-92b *in vitro*, we evaluated the influence of APE1 on miR-92b directly in human cells. Firstly, the ability of APE1 to bind either the precursor form of miR-92b, pri-miR-92b, or its mature form, miR-92b-3p, was investigated by RIP analysis in HeLa clones (70) including: i) SCR, which expresses only the endogenous form of APE1; ii) cl.3, a doxycycline-inducible knock-down (KD) model for APE1 protein, and iii) APE1^WT^ and iv) APE1^NΔ33^ clones, expressing a Flag-tagged ectopic form of APE1 in place of the endogenous ones (70). The input and the FLAG-immunoprecipitated materials, upon RNA extraction procedure, were reverse-transcribed and analyzed by qPCR (Fig. 4A and Fig. S4A). Pri-miR-92b was successfully immunoprecipitated only in the APE1^WT^ clone wheraes, as expected, no signal was obtained in both SCR and cl.3 clones. Furthermore, in accordance with the binding experiments, the APE1^NΔ33^ clone lost about 80% of the immunoprecipitated pri-miR-92b compared to APE1^WT^. We also investigated if APE1 was able to bind miR-92b-3p, the most expressed strand of the two mature forms of pri-miR-92b (Fig. 4B). Noteworthy, miR-92b-3p does not present any rG4-forming motif being cytosine-rich, and indeed, in accordance, RIP analysis showed that the immunoprecipitated materials from APE1^WT^ and APE1^NΔ33^ clones did not present any statistical enrichment of miR-92b-3p when compared to SCR and cl.3 clones.These results indicated that APE1 was able to bind rG4-containing pri-miR-92b also in the cellular context. As pri-miRNAs are prevalently double-stranded, these data would also suggest that the rG4-structure in pri-miR-92b could be in equilibrium with the double-stranded form.

**Figure 4:**
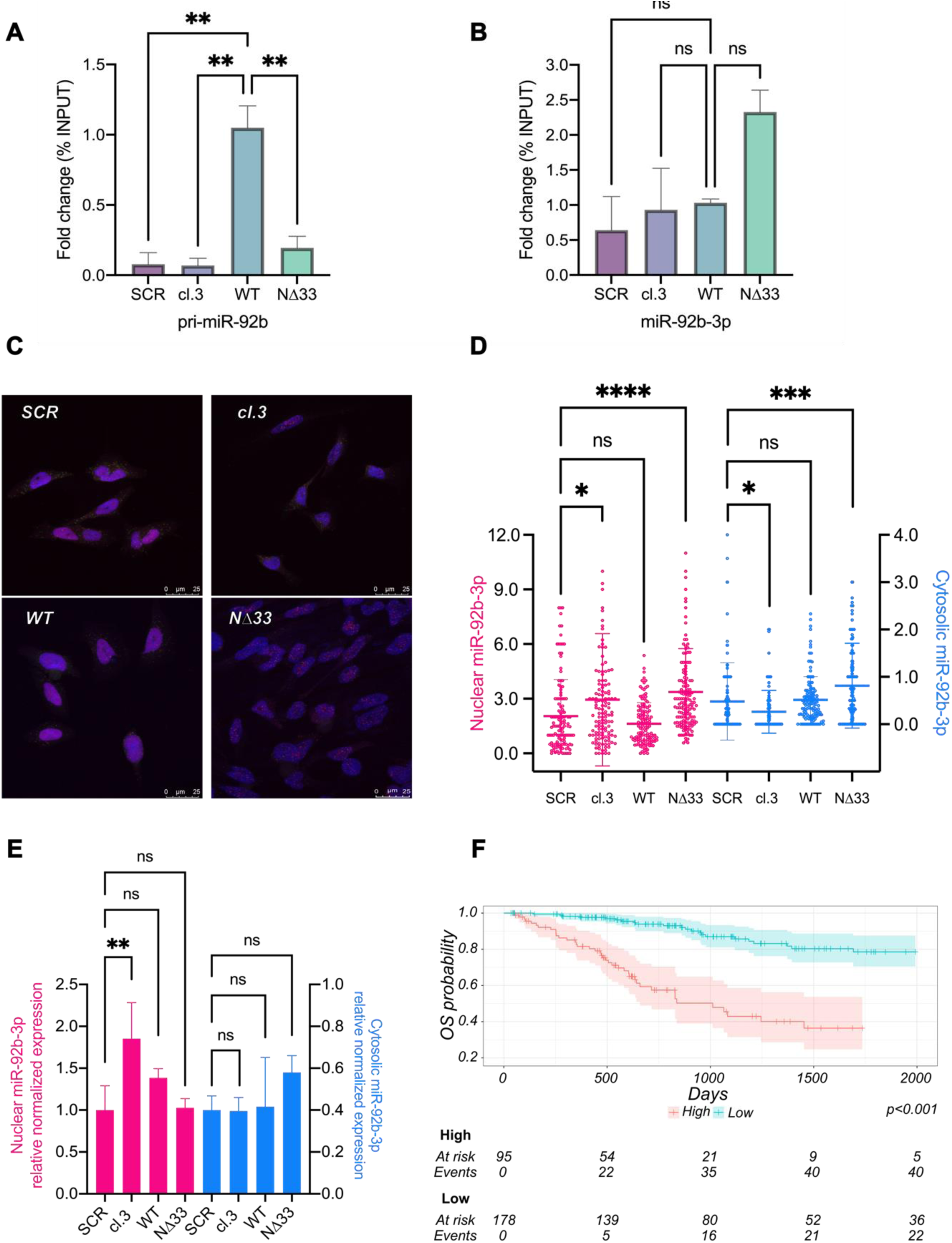
APE1 binds pri-miR-92b in HeLa cells and influences the nucleo-cytosol localization of miR-92b-3p in HeLa cells. A) Validation of APE1 binding towards pri-miR-92b and B) miR-92b-3p in HeLa cell clones by RIP analysis. Data are represented as fold change percentage of the amount of immunoprecipitated target on the amount of the target present in the total input RNA (% INPUT) and normalized to APE1^WT^. The data are expressed as means ± S.D. of two independent replicates. C) Representative fluorescence confocal microscope images of FISH analysis in HeLa clones. Cells were stained with probes specific for miR-92b-3p (red) and actin mRNA (green), as control. APE1 staining was carried out using anti-APE1 specific antibody (magenta) and nuclei were stained by DAPI (blue). D) Scatter plot of FISH analysis representing normalized individual values of miR-92b-3p in the nuclear (in pink) and cytoplasmic (in blue) compartment of HeLa clones silenced for endogenous APE1 expression. Nuclear and cytoplasmic miR-92b-3p dots were counted for each cell, using DAPI as a nuclear mask and APE1 as a cytoplasmic mask. The number of dots of miR-92b-3p was normalized on the number of dots of actin mRNA *per* each cell, in each compartment. The right axis corresponds to the nuclear miR-92b-3p levels, while the left one to the cytoplasmic miR-92b-3p levels. The data are expressed as means ± S.D. of two independent replicates. E) Analysis of miR-92b-3p levels in the nuclear and cytoplasmic compartment in HeLa clones silenced for endogenous APE1 expression by qRT-PCR. The histogram shows the expression levels of miR-92b-3p in the nucleus, normalized on miR-484 and miR-196 levels and the expression levels of miR-92b-3p in the cytoplasmic compartment, normalized to miR-16-5p levels. Data are expressed normalized with respect to SCR. The data are expressed as means ± S.D. of three independent replicates. F) Kaplan–Meier plots in TCGA-CESC dataset show the different OS rates of subjects belonging to the “High risk” and “Low risk” groups, stratified based on the PI calculated from the miRNA signature. Results have been reported from 0 to 2000 days as the sample size after 2000 days is too small to perform statistical inference. A p-value < 0.05 is symbolized by a single asterisk (*), while p-value < 0.001 is symbolized by three asterisks (***) and < 0.0001 by four asterisks (****).

In a previous publication, we demonstrated that cell depletion of APE1 led to an impairment of pri-miRNA processing for some specific miRNAs (41). For this reason, we examined if APE1 was also involved in the maturation of pri-miR-92b. We checked pri-miR-92b and miR-92b-3p expression levels in the above-mentioned HeLa clones by qPCR (Fig. S4B and Fig. S4C). Specifically, pri-miR-92b levels were almost comparable between SCR and cl.3, with a slight decrease (15%) in the APE1-depleted HeLa clone, in line with previously described (41, 43). Furthermore, when APE1 expression was rescued by expression of siRNA-resistant ectopic APE1^WT^ or APE1^NΔ33^ proteins (70), pri-miR-92b levels resulted comparable to the SCR. Moreover, we did not observe any significant changes in the levels of miR-92b-3p among the different clones (Fig. S4D).

These data indicate that APE1 is able to bind to the miR-92b precursor form, whereas its overall expression is only slightly influenced by APE1 depletion.

Recently, an important role of miRNA subcellular distribution in their regulation has been pointed out (71), thus we speculated that APE1 could influence and regulate the distribution of miRNAs between nuclei and cytosol. First, we evaluated the impact of APE1 depletion in the subcellular localization of miR-92b-3p by quantification through FISH analysis in HeLa clones (Fig. 4C, Fig. S5A and Fig. S5B). In basal conditions (SCR), miR-92b-3p localization was both nuclear and cytoplasmatic (red dots), being more present in the nucleus. Upon APE1-knockdown (cl.3), miR-92b-3p was enriched in the nuclear compartment with the amount of nuclear miRNA significantly increased by about 43% compared to SCR, while its levels were significantly decreased in the cytosol by about 46% (Fig. 4D). When APE1 expression was rescued by the ectopic APE1^WT^-Flag protein, the localization was comparable to SCR. Contrarily in the APE1^NΔ33^ rescue, we observed a higher number of red dots compared to SCR and WT clones, in both nuclear and cytoplasmic compartments, probably because APE1^NΔ33^ lacks its nuclear localization signal and is located in the cytosolic compartment. We confirmed FISH data by performing quantitative measurements through qPCR analyses on RNA obtained from nuclear and cytoplasmic compartment subfractionation experiments. Thus, we isolated RNA and proteins from cytoplasmic and nuclear cell fractions from HeLa clones. First, we confirmed the efficiency of fractionation by performing a WB analysis (Fig. S4E). Lamin A antibody was used as a control for the proper isolation of nuclei, while tubulin and Hsp70 for the cytosol. APE1 resulted prominently nuclear in SCR and APE1^WT^ clones, while predominantly cytoplasmic in APE1^NΔ33^ clones, as expected. Afterward, we analyzed the levels of miR-92b-3p in the nuclear and cytoplasmic compartments by qPCR. In Fig. 4E, we showed that the APE1 depletion led to an accumulation of miR-92b-3p in the nucleus with a one-fold increase compared to SCR. Remarkably, the amount of cytoplasmic miR-92b-3p slightly decreased in APE1-depleted cells. Following the ectopic expression of APE1^WT^-Flag, the localization of miR-92b-3p resulted almost comparable to what observed in SCR. Finally, the rescue with the ectopic APE1^NΔ33^-Flag protein showed a similar trend of enrichment in the nucleus and the cytosol supporting the FISH results, although not statistically significant.

Overall, these data suggested that APE1 is involved in the maturation process and shuttling between the nucleus and cytoplasm of miR-92b, thus regulating its amount in both compartments.

Lastly, to explore the prognostic power of rG4-miRNAs regulated by APE1 in different cancer types, we retrieved both miRNA expression and clinical data of TCGA samples for Cervical Squamous Cell Carcinoma and Endocervical Adenocarcinoma (TCGA-CESC), Lung Adenocarcinoma (TCGA-LUAD), and Liver Hepatocellular Carcinoma (TCGA-LIHC). These specific tumour types were chosen to be consistent with the cellular models used in this work and previous studies of our laboratory. For each dataset, we computed a Prognostic Index (PI) to stratify subjects into “High risk” and “Low risk” groups, based on both miRNA expression and association with overall survival (OS). In particular, the subject’s PI was computed using multivariate Cox regression coefficients and the expression values of selected miRNAs for which expression data are available in the corresponding TCGA dataset (46 miRNAs in TCGA-LUAD and TCGA-CESC and 16 miRNAs in TCGA-LIHC). Subjects were then stratified into the two groups, based on p-value optimization. Finally, Kaplan-Meier curves and a log-rank test were used to summarize data and compute the associated p-value. The OS rates were significantly different between the “High risk” and “Low risk” groups in all the analysed datasets (TCGA-CESC, Fig. 4F, High risk = 95 and Low risk = 178, p-value = 5 x 10^-14^, Hazard ratio Low risk / High risk = 0.17, TCGA-LUAD, Fig. S6A, High risk = 142 and Low risk = 285, p-value = 1 x 10^-12^, Hazard ratio Low risk / High risk = 0.33, TCGA-LIHC, Fig. S6B High risk = 118 and Low risk = 221, p-value = 3 x 10^-8^, Hazard ratio Low risk / High risk = 0.37) suggesting that the expression of the miRNA signature is significantly associated with a different OS probability and supporting their potential prognostic value in the three analysed cancer types. To identify the minimum miRNA signature associated with changes in the survival rate and to prioritize miRNAs based on their contribution, we then fitted a Cox regression model with LASSO penalization. The resulting LASSO lambda coefficients were used to rank miRNA based on their importance. In this case, miRNAs showing a coefficient equal to 0 have little to no impact on the signature and were thus excluded from the minimum miRNA signature. In particular, the minimum miRNA signature is composed of the following miRNAs in the three datasets analysed: in TCGA-CESC mir-642a, mir-1275, mir-1250, mir-1249, mir-296, mir-541, mir-320a, mir-1292, mir-1226 and mir-328 (Fig. S6C), in TCGA-LUAD mir-1246, mir-3911, mir-320d-2, mir-92b, mir-328, mir-3940, mir-584, mir-1292, mir-940, mir-3195, mir-3187, mir-134, mir-1250, mir-541, mir-185, mir-642a, mir-200c, mir-615, mir-636, mir-9-1, mir-3928, mir-3615, mir-671, mir-877, mir-139b and mir-675 (Fig. S6E) and in TCGA-LIHC mir-1249, mir-3127, mir-197, mir-139, mir-1226 and mir-1250 (Fig. S6G). Finally, to confirm the prognostic power of the minimum miRNA signature, we computed the PI using the LASSO lambda coefficients instead of the Cox coefficients. In this way, only miRNAs having a lambda coefficient different from 0 are considered. Subjects were then stratified into the “High risk” and “Low risk” groups based on p-value optimization, and a Kaplan-Meier curve along with a log-rank test was used to test the difference in the OS probability between the two groups. Interestingly, the minimum miRNA signature was still able to explain the difference in the OS probability between the two groups in all the analysed datasets (TCGA-CESC, Fig. S6D, High risk = 122 and Low risk = 151, p-value = 6 x 10^-8^, Hazard ratio Low risk / High risk = 0.27, TCGA-LUAD, Fig. S6F, High risk = 159 and Low risk = 268, p-value = 9 x 10^-13^, Hazard ratio Low risk / High risk = 0.32, TCGA-LIHC, Fig. S6H, High risk = 114 and Low risk = 225, p-value = 4 x 10^-11^, Hazard ratio Low risk / High risk = 0.30).

These results, together with additional analysis on their target genes (Fig. S7), supported the hypothesis that a major role of APE1 in tumour progression could be ascribable to its regulation of rG4-containing onco-miRNA sequences.

## Discussion

rG4 structures are post-transcriptional regulators of gene expression (72). Their folding and shuttling are controlled by RNA binding proteins (i.e. hnRNPA2B1, FUS, NCL, NPM1 etc.), RNA helicases (i.e. DHX36, hnRNP H/F (73, 74)), cations (i.e. Na^+^ and K^+^) and small molecules that are usually planar chromophores such as fused aromatic polycyclic systems, macrocycles and non-fused aromatic systems with flexible structural motifs (10), making them highly dynamic (75). rG4s are particularly enriched in miRNAs (76). Recent data underline an emerging regulatory function played by rG4s in miRNA-maturation through Drosha and Dicer inhibition and a potential role in physiological and pathological liquid-liquid phase separation mechanisms (77, 78). In addition, several evidence have shown that the presence of the rG4s in pre-miRNA is in equilibrium with the canonical stem-loop structure, and may regulate the maturation of some miRNAs, such as miR-63b (73), miR-1587 (22, 79), miR-26a-1 (18), miR-1229 (19), miR-3620-5p (24), pre-let-7 (80), and miR-92b (28, 33), thus offering a new layer of gene expression regulation through post-transcriptional mechanisms. Since dysregulation of miRNA biogenesis, as well as the quality control mechanisms of damaged miRNAs processes, are emerging as hallmarks of cancer progression, it is essential to understand the regulatory aspects of these processes (81). In particular, defining the regulatory role of non-canonical structural elements, including rG4s, on miRNA function, maturation, shuttling, and processing would represent a step forward in translating these mechanisms into therapeutic opportunities for personalized medicine.

In literature, it has been highlighted the importance of APE1 in gene expression regulation, also mediated by G4 structures in DNA (36, 40, 82). Indeed, through its redox domain, APE1 activates several TFs (48), but it preferentially binds to G4 of several gene promoters, such KRAS (36), NEIL3, VEGF (40, 82), and induces the formation of transcription hubs, enhancing gene expression. APE1 also regulates gene expression through post-transcriptional mechanisms involving miRNAs (35, 41, 48, 83). One example is represented by the expression of PTEN tumor suppressor, which is regulated by APE1 by two complementary mechanisms (41, 84). Indeed, APE1 activates the Egr-1 transcription factor through its redox domain, enhancing PTEN gene expression (84), while it also regulates miR-221/222 maturation, which post-transcriptionally targets PTEN, leading to its mRNA degradation (41). Given all these roles of APE1, the present study was conceived and designed to address the involvement of APE1 in the regulation of rG4-containing miRNAs.

In recent years, we aimed at the identification of those miRNAs whose expression is regulated by APE1, using different cancer cell lines (41, 43, 83). Surprisingly, between these miRNAs, we found an enrichment of those bearing an rG4 motif in their immature forms (Fig. 1A). We employed miR-92b as a model, for the reported role of the rG4 structure present in its precursor form (pre-miR-92b) in regulating its maturation (28, 33), and as it targets PTEN, whose expression, in turn, is regulated by APE1 at the transcriptional and post-transcriptional levels (41, 84). Here, we demonstrated that APE1 is able to bind and dynamically regulate the rG4 structure in the pre-miR-92b. The biochemical characterization showed that the affinity of this binding is in the low micromolar range, which is in agreement with the biological relevance of this interaction. Moreover, structural analyses through CD and NMR confirmed that the pre-92b adopts a parallel-stranded G4 in solutions with K^+^ ions.

Of interest, we observed that the 33 N-terminal region of APE1 and, in particular, the lysine residues at position 27/31/32/35, are directly involved in the stabilization of the binding between APE1 and the rG4-containing pre-miR-92b. These findings are in agreement with our previous evidence, underlining the relevance of these lysine residues in modulating the activity of APE1 on different RNA molecules (37, 39, 85). In addition, we showed that important flexible residues such as the RM bridge and other residues located in the catalytic domain and the hydrophobic product pocket, are key players in the interaction with pre-92b probe (Fig. S3).

Noticeably, FISH and qPCR cellular assays pointed out that APE1 cellular depletion causes miR-92b retention in the nucleus (Fig. 4C, 4D and 4E) and these data are compatible with the general hypothesis that APE1, through its ability to bind and modify rG4 structures present in the precursor form of miR-92b, is involved in the maturation/shuttling process of the mature miR-92b, thus possibly contributing to miRNA biology and the post-transcriptional control of gene expression (71, 86, 87). Whether APE1 may act as a ‘simple chaperone’ for rG4-containing miRNAs or take part in the enzymatic process involved in their maturation is currently unknown and should be further investigated.

Intriguingly, we observed that genes targeted by the APE1-regulated rG4-containing miRNAs are associated with several pathways related to cancer onset and progression in the LUAD, CESC, and LIHC datasets. Besides canonical cancer-related pathways like “small cell lung cancer” and “hepatocellular carcinoma”, we also highlighted more specific pathways related to cell proliferation and apoptosis. Finally, it is also worth mentioning that pathways related to cell-to-cell and cell-to-extracellular matrix interactions are enriched with APE1-regulated miRNA genes suggesting a role of APE1 in the epithelial–mesenchymal transition.

As previously described in our and others literature (48, 88–91), APE1 is overexpressed and serum-secreted in hepatocellular (HCC), non-small cell lung (NSCLC), and colon (CRC) cancers, representing a poor-outcome prognostic/predictive factor and an emerging anti-cancer target. Moreover, APE1 regulates oncogenic miRNA maturation and decay of oxidized- and abasic-miRNAs and can be secreted by cancer cells through extracellular vesicles (41, 88–91). By analyzing the TCGA datasets, we showed that the expression profile of rG4-containing miRNAs regulated by APE1 represents a promising tool to define a signature that clearly separates patients into two distinct groups characterized by a significant difference in the OS probability. This suggested the existence of a relationship between miRNA expression and survival in patients affected by lung adenocarcinoma, cervical cancer and hepatocellular carcinoma and confirmed the important prognostic value of those miRNAs in the analyzed cancer types (Fig. 4F, Fig. S6 and Fig. S7).

As recently proposed (92), after genotoxic treatment, damaged miRNAs including those containing AP or 8-oxoguanine sites, seem to accumulate up to 10-fold or more in cells, having fundamental effects on gene expression (93), and an important epitranscriptional regulation on cancer development. Whereas in duplex DNA the electron holes can travel over long distances (from 100s to 1’000s bp) until reaching a GGG site (as found in a G4 motif) with a low redox potential site thus forming an 8-oxoguanine, in the case of RNA, this does not happen. Currently, the frequency and extent of AP and 8-oxoguanine sites in miRNAs is still unknown. It is plausible that guanine oxidation may rewire miRNAs by differentially regulating redox-dependent cancer development as well as the modulation of miRNA-processing and decay during genotoxic stress may be part of the executive mechanism of chemoresistance (92, 94). Not least, the biochemical mechanisms responsible for the repair of oxidized bases and the generation of AP sites in miRNAs remain unknown. On these topics, the ability of APE1 to recognize and process miRNAs containing both AP and 8-oxoguanine sites, alone or in combination with additional proteins such as hnRNPD (95) or PCBP1 (96), may represent an interesting subject of investigation focused on defining the regulatory functions of rG4s present in several miRNAs (92, 93), and for understanding molecular mechanisms in miRNAs quality control, with the ultimate goal of developing novel specific anticancer strategies. Future work in our laboratory will address these issues.

### Material and Methods

#### Bioinformatic rG4 prediction, functional enrichment analysis and network miRNA-gene generation

For further detailed information, refer to SI Appendix.

#### Synthetic oligoribonucleotides, description, and annealing conditions

For further detailed information, refer to SI Appendix.

#### Expression of recombinant proteins and FPLC purification

Expression of human recombinant GST-tagged and untagged APE1^WT^, APE1^NΔ33,^ and APE1^K4pleA^ proteins was obtained as explained in (39).

#### SDS-PAGE and Western Blotting (WB) analysis

SDS-PAGE and WB were conducted as explained in (41). The primary antibodies used and their dilution usage were following listed: APE1, mouse, monoclonal, 1:2000; Novus; Actin, rabbit, polyclonal, 1:2000, Sigma; β-tubulin, mouse, monoclonal, 1:2000, Sigma; lamin-A, mouse, monoclonal, 1:1000, abcam; Hsp-70, rabbit, polyclonal, 1:10’000, GeneTex.

#### Northwestern Blot Assay (NWB)

NWB assay was performed as explained in (95), and incubating with 5 pmol of the labeled oligoribonucleotide. After washing, the membranes were scan by Odyssey CLx scanner and analyzed byImageStudio Software (Li-Cor Biosciences).

#### RNA Electrophoretic mobility shift assay analysis (REMSA)

For the REMSA analysis, the reactions were prepared in a final volume of 20 μl with the indicated doses of APE1 recombinant protein and 25 nM of the oligoribonucleotide, in a buffer containing 20 mM TrisHCl pH 7.4, 100 mM KCl, 5 µg/µl BSA and 0.25 % glycerol. The reactions were incubated for 30 minutes at 4°C and then added with 2 μl of Orange Loading Dye (Li-Cor Biosciences). The samples were then run on a 4 % native PAGE gel at 60 V for the first 15 minutes and then increasing at 80 V for the remaining 60 minutes, at 4°C in TBE 0.5 X. After running, the gels were scan by Odyssey CLx scanner and analyzed byImageStudio Software (Li-Cor Biosciences).

#### Binding assay by UV-crosslinking

For the UV-crosslinking analysis, the reactions were prepared in a final volume of 20 μl with the indicated doses of APE1 recombinant protein and 25 nM of the oligoribonucleotide, in a buffer containing 20 mM TrisHCl pH 7.4, 100 mM KCl, 0.5 µg/µl BSA and 0.25 % glycerol. The reactions were incubated for 30 minutes at 4°C and then UV-crosslinked at 0.2 J/m^2^. The samples were then added with laemmli 4X and heated at 95°C for 5 minutes. The samples were run on a 12 % SDS-PAGE gel. After running, the gels were scan by Odyssey CLx scanner and analyzed by ImageStudio Software (Li-Cor Biosciences).

#### MicroScale Thermophoresis (MST) experiments

MST experiments were carried out with a Monolith NT 115 system (Nano Temper Technologies) equipped with 20% LED and 40% IR-laser power. For the assay, a 16-step serial dilution (1:1) procedure was performed, with the final concentrations of 12.5 μM for APE1^WT^, 13.5 μM for APE1^NΔ33^ and 5 μM for APE1^K4pleA^; pre-92b probe was added in each tube at a final concentration of 625 nM. The samples were filled into standard capillaries and measurements were carried out at 25°C in a buffer containing 25 mM Tris-HCl pH 7.5, 100 mM NaCl, 1 mM DTT. The equation, used for fitting data at different concentrations was implemented by the software Graph PAD with the equation nonlinear regression log (inhibitor) vs. response (three parameters).

#### Circular Dichroism (CD) analysis and UV and fluorescence spectroscopies

For further detailed information, refer to SI Appendix.

#### Cell culture and compounds

HeLa cells were grown as indicated in (41). HeLa stable clones were grown as indicated in (70) and harvested 10 days after the addition of doxycycline in the medium.

#### RNA extraction and quantitative Real Time PCR (qRT-PCR)

For the separation of nuclear and cytosolic RNA, we followed what was described in (97). Total, nuclear and cytosolic RNA pellets were resuspended in Qiazol and RNA was extracted using miRNeasy kit (Qiagen, USA), according to the manufacturer’s instructions. For qRT-PCR information, refer to SI Appendix.

#### RNA Immunoprecipitation (RIP) analysis

RIP analysis on HeLa cell clones was performed as previously described in (41).

#### Fluorescence In Situ Hybridization (FISH) analysis

Doxy-treated HeLa cell clones were plated on coverslips in 24-multiwell and grown to 60-80% confluency. Coverslips were stained using ViewRNA^TM^ Cell Plus Assay Kit (Invitrogen, ThermoFisher), according to the manufacturer instructions. The detection of the targets was carried out using miR-92b-3p Type 1 probe (VM1-10117-01, ThermoFisher), miR-16-5p Type 1 probe (VM1-10232-01, ThermoFisher) and ACTB, human Type 4 probe (VA4-10293-01, ThermoFisher). APE1 was stained by using anti-APE1 antibody (mouse, monoclonal, 1:100, Novus) and Alexa® 633 secondary antibody (Invitrogen, A211050). All images were captured by confocal microscopy (Leica) in z-stack. Maximum projections were analyzed using ImageJ software, as described in (98).

#### NMR titration

Both 1D and 2D NMR experiments were recorded on a Bruker Advance III 700 and 800 MHz spectrometer equipped with a liquid TXI 1H/13C/15N/2H probe. Samples (3 mm NMR tubes) were prepared in Kpi buffer 10 mM K_2_HPO_4_/KH_2_PO_4_; 50 mM KCl, 2 mM MgCl_2_, 0.2 mM TCEP at pH 6.8. D_2_O (8 %) was used as reference. APE1 was used at 100 (1D) or 64 (2D) µM. In the 1D (1H) NMR experiments, the water signal was suppressed using excitation sculpting with gradients (zgesgppe; d1=2sec; 512 scans; time domain=64k). We used Transverse Relaxation-Optimized Spectroscopy (TROSY) to acquire the 2D titration spectra. Each residue has been identified by the −NH from its backbone connection and assigned using the deposited data from PDB structure 1BIX and 1DEW together with data available at the BMRB entry 16516. NMR data was treated using TopSpin 4.1, NMRFAM-Sparky (99), (Bruker Biospin), and the structure analysis was performed UCSF Chimera (100).

#### Survival analysis

For further detailed information, refer to SI Appendix.

## Supporting information

Supplementary Informations

## Acknowledgments

The work was supported by: Associazione Italiana per la Ricerca sul Cancro (AIRC) with [grant number IG19862] to G. T. and [grant IG27378] to D.M., through the support of the Departmental Strategic Plan (PSD) of the University of Udine-Interdepartmental Project on Artificial Intelligence (2020–25), by additional grants from the University of Udine (‘Bando Ricerca Collaborativa’ granted by European Community - NextGenerationEU) and from the Consorzio Interuniversitario Biotecnologie - C.I.B. – (MUR-PRIN2022: “L’INNOVAZIONE DELLE BIOTECNOLOGIE NELL’ERA DELLA MEDICINA DI PRECISIONE, DEI CAMBIAMENTI CLIMATICI E DELL’ECONOMIA CIRCOLARE”) to G.T.. Moreover, C.J.B. and A.F.M. thank the U.S. National Institutes of Health via R35 GM145237 for financial support. Finally, G.S. thanks the IR INFRANALYTICS FR2054 and the IECB NMR platform at the University of Bordeaux.

