## Supplementary Informations for "The Apurinic/Apyrimidinic Endodeoxyribonuclease 1 is an RNA G-quadruplex binding protein and regulates miR-92b expression in cancer cells"

**This PDF file includes:**

Supporting Text
Supporting Methods

Figures S1 to S7
Table S1
Legends for Dataset S1, Dataset S2
References

**Other supporting materials for this manuscript include the following:**

Dataset S1
Dataset S2

### Supporting Text

**APE1 depletion affects the expression of genes involved in different cancer-related processes**

To shed light on the biological processes associated with genes targeted by the APE1-regulated rG4-containing miRNAs, we performed an enrichment analysis. To select only context-meaning target genes, we included some additional filters. In particular, we retained only miRNA target genes that were differentially expressed upon APE1 *KD* in HeLa cells (1) and whose expressions anticorrelate with that of their regulatory miRNAs resulting in 156 in CESC, 209 target genes in LUAD and 119 in LIHC. The results of the enrichment analysis are reported as bar plots representing the most important enriched pathways in each dataset, sorted by adjusted p-value, along with a gene-pathway network that also shows some of the most important genes in each pathway and highlights shared genes among different pathways (Fig. S6 and Fig. S7). Interestingly the results obtained were rather concordant among all datasets analysed, reporting several common pathways. We observed different enriched tumorigenic pathways, such as “p53 signalling” (adj p-value LIHC = 0.03), and pathways associated to specific tumour types such as “small cell lung cancer” (adj p-value LUAD = 0.0001, CESC = 0.007), “gastric cancer” (adj p-value CESC = 0.04), “hepatocellular carcinoma” (adj p-value CESC = 0.026) and “colorectal cancer” (adj p-value LUAD = 0.04, CESC = 0.006), further confirming the relevance of this miRNA signature in cancer onset and progression in the analysed datasets. Furthermore, the analysis also highlighted more specific pathways involved in cancer progression. The “PI3K-AKT signalling pathway” (adj p-value LUAD = 0.005) contained important oncogenic genes such as PTEN that we previously demonstrated to be regulated by APE1. We also observed enriched pathways related to tumour suppression like “Hippo signaling pathway” (adj p-value LUAD = 0.04).and “FoxO signalling” (adj p-value CESC = 0.02), whose deregulation may imbalance some important cancer-related cellular processes like as cell proliferation, cell growth and apoptosis (2–5). Moreover, pathways related to the cell cycle and some immune-related pathways like “Human T-cell leukaemia virus infection” (adj p-value CESC = 0.04), “Human cytomegalovirus infection” (adj p-value LUAD = 0.04, CESC = 0.04), “Viral carcinogenesis” (adj p-value CESC = 0.02, LUAD = 0.04) and “Human papillomavirus infection” (adj p-value CESC = 0.04) were found to be enriched of miRNA targets genes extending the relevance of the miRNA signature also to immune-related cancer pathways (6). We also highlighted some other important pathways related to cell-to-cell and cell-to-extracellular matrix interaction (“ECM-receptor interaction” (adj p-value CESC = 0.04), “Proteoglycans in cancer” (adj p-value CESC = 0.003, LUAD = 0.005, LIHC = 0.01) and “Focal adhesion” (adj p-value LUAD = 0.002, CESC =0.003) that might suggest a relation with processes such as cell migration, invasion, and epithelial-to-mesenchymal transition. These pathways play pivotal roles in cancer onset and progression and had previously been associated with APE1 overexpression (7–11). Finally, the “micro RNAs in cancer” (adj p-value CESC = 0.05, LUAD = 0.04) pathways were found to be significantly enriched, confirming the role of APE1 in cancer-relevant miRNA biogenesis (7). Taken together, the enrichment of these pathways confirmed a strong association between APE1-regulated rG4-containing miRNAs and tumor onset and progression.

### Supplementary Methods

**Bioinformatic rG4 prediction, functional enrichment analysis and network miRNA-gene generation**

Pre-miRNA and mature miRNA sequences were extracted as FASTA files from human miRNA sequences download miRbase (12). rG4s were then predicted on these sequences by running the QGRSmapper (score threshold = 19, (13)), psqfinder (score threshold = 47, (14, 15)) and G4Hunter (parameters -w 25 -s 1.2, (16, 17)) algorithms. Finally, the obtained rG4s were intersected to extract those predicted by one, two, or all three algorithms. Experimentally identified rG4s were also searched at the miRNA gene level by using the Batch Download function of the QUADRatlas database (18). To identify deregulated pathways associated with APEX1 KD, we have performed an enrichment analysis using ClusterProfiler by querying the KEGG database on the union of upregulated genes from HeLa and A549 cells (q-value < 0.05). In order to decipher the regulatory miRNA-gene network of the 61 miRNA signature, we retrieved the associated target genes from Tarbase. Target genes were then filtered to maintain only upregulated genes in both HeLa and A549 APEX1 KD cells. Upregulated genes were defined, for both cell lines, as genes having a q-value < 0.05 and a log2FoldChange > 0. The miRNA-gene network has been constructed using the igraph R package.

**Synthetic oligoribonucleotides, description, and annealing conditions**

For NWB, REMSA, UV-crosslinking and MST analysis, we used a twenty-nucleotide long oligoribonucleotide with the following sequence 5’-CCGGGCGGGCGGGAGGGACG-3’, indicated as pre-92b. The oligoribonucleotide hold an IRDye-700 at the 5’ and was synthesized by Metabion, purified by HPLC and checked in Mass Spectrometry. It was resuspended in DNase- and RNase- free water, at a final concentration of 100 µM. Annealing was performed at a final concentration of 2.5 µM in a solution containing either 100 mM KCl or 100 mM LiCl, heated at 87°C and cooled down overnight. For CD analysis and fluorescence spectroscopy, the pre-92b probe was also synthesized (5′-CCGGGCGGGCGGGAGGGAC-3’) with and without a FAM fluorophore at the 3’ end, by solid-phase synthesis using standard protocols. The complement, holding a black hole quencher (BHQ) at the 5’ (5′-BHQ-GGCCUCCGGCCCCCCCG-3’) was also synthetized by solid-phase synthesis using standard protocols. The strands were purified by anion-exchange HPLC as previously described (19). The oligonucleotide was annealed at 5 µM concentration in 20 mM Tris pH 7.4 (at 22 °C) and 50 mM potassium acetate by heating it to 90 °C for 5 min in a water bath followed by slowly cooling it to room temperature.

**Circular Dichroism (CD) analysis and UV and fluorescence spectroscopies**

The oligoribonucleotide sample was placed in 0.2-cm quartz cuvette for CD analysis at 22 °C. The CD spectrum was recorded from 220-320 nm, followed by background subtraction of the solvent signal, and then the data were normalized to molar ellipticity values for visualization. The thermal melting value for a 2-µM concentration of the rG4 was measured as previously described (19). The duplex to rG4 APE1 induced structural conversions were determined on the 3′-FAM labeled rG4 annealed to the 5′-BHQ labeled complement as described above. The fluorescence intensity at 520 nm was monitored under the same conditions for the binding studies on a 100 nM RNA solution that was titrated with a 2-fold serial dilution of the protein starting at 5 µM. The $FI_{520nm}$ values were normalized and then processed as previously described (19).

**Thermal denaturation**

Thermal denaturation profiles were evaluated by recoding CD spectra on a Jasco J-810 spectropolarimeter (JASCO Corp., Easton, MD) equipped with a Peltier temperature controller in a 220−260 nm interval with 5 μM protein concentration in 5 mM phosphate buffer (pH 7.5) and 250 mM NaCl with a 0.1 cm path cuvette. Thermal denaturation profiles were obtained by measuring the temperature dependence of the signal at 230 nm in the range of 20−80°C with a resolution of 0.5°C and a 1.0 nm bandwidth. Data were collected at 0.2 nm resolution with a 20 nm/minute scan speed and a 4 s response and were reported as the unfolded fraction versus temperature.

**RNA extraction and quantitative Real Time PCR (qRT-PCR)**

Hsa-miR-92b-3p and hsa-miR-92b-5p were validated by qRT-PCR using TaqMan Advanced miRNA assay (Life Technologies, USA), as indicated by the manufacturer's instructions. The detection of the retro-transcribed products was obtained using TaqMan Fast Advanced Master Mix and CFX Touch Real-Time PCR system (Bio-Rad). The results of qRT-PCR were obtained using the $\Delta\Delta C_t$ method, normalizing for the expression of miR-16-5p as a reference for total and cytosolic fractions, while normalizing for miR-196b and miR-484 for nuclear factions. For the measurement of pri-miRNA levels, one microgram of RNA was reverse transcribed with SensiFAST cDNA synthesis kit (Bioline, UK), following the manufacturer's instruction. The detection of the retro-transcribed products was obtained using Taqman pri-miRNA probe and TaqMan Fast Advanced Master Mix and CFX Touch Real-Time PCR system (Bio-Rad). The results of qRT-PCR were obtained using the $\Delta\Delta C_t$ method, using the expression of GAPDH as a reference.

**Survival analysis**

The prognostic value of selected miRNAs was evaluated in three datasets retrieved from The Cancer Genome Atlas (TCGA) including Lung Adenocarcinoma (TCGA-LUAD), Cervical Squamous Cell Carcinoma, and Endocervical Adenocarcinoma (TCGA-CESC), and Liver Hepatocellular Carcinoma (TCGA-LIHC). miRNA expression for tumor samples (Illumina HiSeq platform, miRgene level, Normalized, RPM), and clinical data were downloaded from the LinkedOmics portal for TCGA-LUAD, TCGA-CESC, and TCGA-LIHC (http://linkedomics.org/data_download/TCGA-LUAD/, http://linkedomics.org/data_download/TCGA-CESC/, http://linkedomics.org/data_download/TCGA-LIHC/; last accessed: June 26, 2023). To evaluate the global prognostic value of the miRNA signature, Cox coefficients were defined for each miRNA based on the multivariate Cox proportional hazard model built using miRNA expression as covariates. These coefficients were then multiplied by the expression value of the associated miRNA for every subject, resulting in a miRNA score. The Prognostic Index (PI) was defined as the sum of all miRNA scores for each subject. Subjects were stratified into "High-risk" and "Low-risk" groups based on p-value optimization using the "*surv_cutpoint*" function from the R "survival" package, with a minimum proportion of 0.33. The difference in overall survival rates between the two groups was assessed using the log-rank test, and a Kaplan-Meier curve was plotted to summarize the data. For the biological pathways associated with deregulated miRNAs target genes, we have performed an enrichment analysis using ClusterProfiler by querying the KEGG database (adjusted p-value < 0.05). Human validated miRNA target genes were retrieved from Tarbase. miRNA target genes were then filtered to maintain only genes that were differentially expressed upon APEX1 KD in the HeLa cell line (*q-value* < 0.05) and that whose expression anticorrelates with the expression of their regulatory miRNA ( LUAD n = 209, CESC n = 156, LIHC n = 119).

**Minimum miRNA signature identification and survival analysis**

To identify the minimum miRNA signature capable of explaining the difference between the two groups of subjects and to prioritize miRNA based on their contribution to the signature a multivariate Cox model with LASSO penalization (alpha = 1) was fitted. The obtained lambda coefficients for each miRNA were used to rank miRNA contribution based on their importance and a PI score was calculated using lambda coefficients instead of the Cox coefficients (miRNA with lambda coefficients equal to 0 were classified as less important features and do not have any impact on PI computation). Finally, to confirm that the minimum miRNA signature is still able to explain the difference in the overall survival probability subjects were stratified into "High risk" and "Low risk" groups, based on *p-value* optimization, and a Kaplan-Meier curve along with a log-rank test was performed.

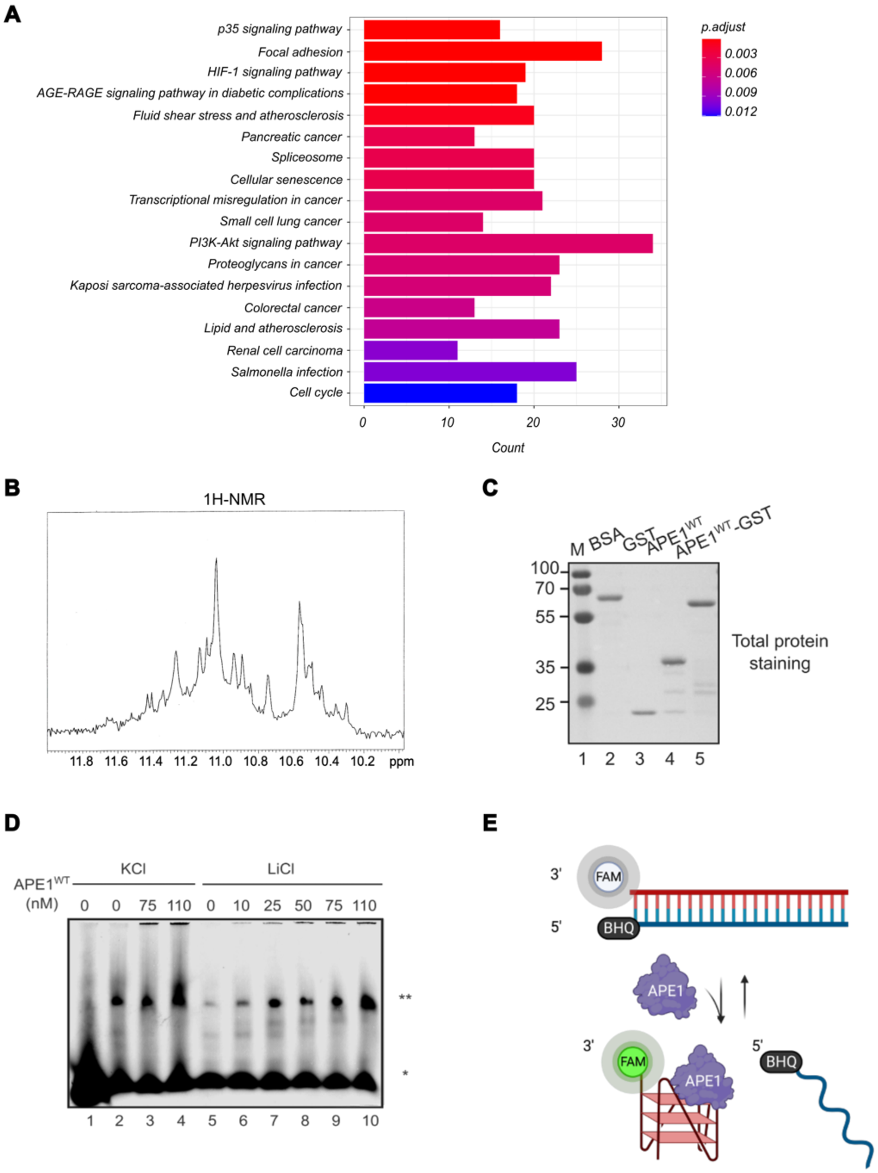


**Fig. S1.** A) Barplot of top 18 significantly enriched pathways, sorted by adjusted p-value, for the union of upregulated genes in HeLa and A549 APEX1 KD cell lines. The x-axis reports the number of upregulated genes in each pathway while the color of the bars represents the associated adjusted p-value. B) NMR spectrum of pre-92b oligonucleotide. C) Total protein staining showing the loading of equal amounts of the recombinant proteins BSA, GST, APE1$^{WT,}$ and APE$^{WT-}$GST. The membrane was colored with Ponceau Red S and acquired with FireReader V10, Uvitec. On the left, the electrophoretic marker is loaded, and the different molecular weights are expressed in kDa. D) Representative REMSA 4% polyacrylamide gel with different quantities of recombinant APE1$^{WT}$ (reported upon the gel and expressed in nM), incubated with a constant concentration of pre-92b probe (25 nM) in 100 mM KCl and 100 mM LiCl, respectively. The asterisks (*), (**) indicate the free probe band, the oligomer structure band. E) Schematic representation of fluorescence anisotropy assay between APE1 and pre-92b probe to assess its structural conversion from duplex RNA to rG4. A 3′-FAM-labeled pre-92b probe was annealed with a complement bearing a 5′ quencher (BHQ), and the sequence provided the two natural bulges predicted in the pre-mir-92b structure. The protein was titrated into the RNA solution while monitoring the evolution of the FAM fluorescence as the structure switched from duplex to rG4. (Created with biorender.com

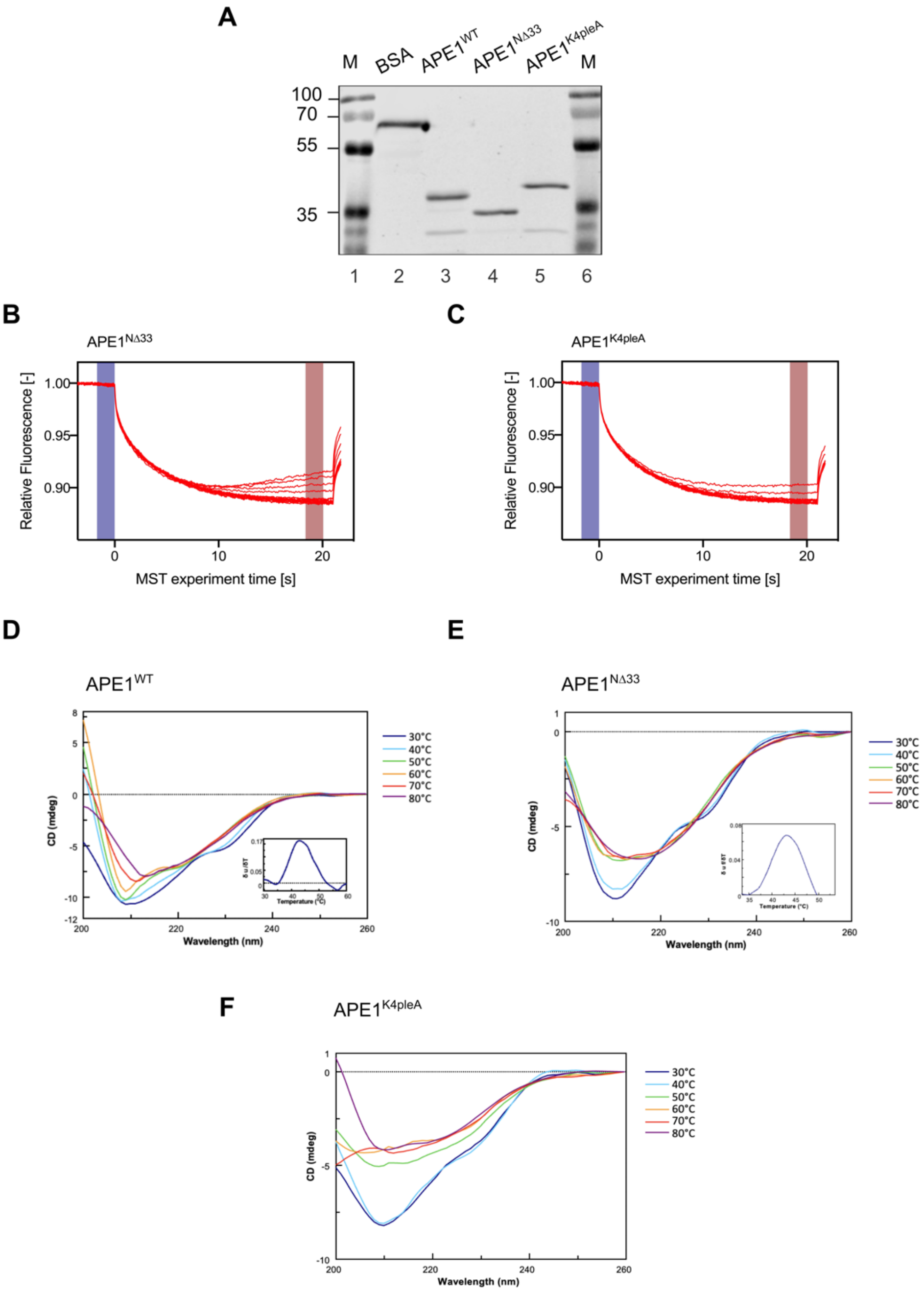


**Fig. S2.** A) Total protein staining showing the loading of equal amounts of the recombinant proteins BSA, APE1$^{WT}$, APE1$^{N\Delta33}$, and APE1$^{K4pleA}$. The membrane was colored with Revert 700 Staining and acquired with Odyssey CLx scanner/ImageStudio Software (Li-Cor Biosciences). On the sides, the electrophoretic marker is loaded, and the different molecular weights are expressed in kDa. B-C) MST between pre-92b and APE1$^{N\Delta33}$ (B) and APE1$^{K4pleA}$ (C). D-F) Overlay of CD spectra registered at indicated temperatures, as inset first derivative of denaturation curves, for APE1$^{WT}$ (D), APE1$^{N\Delta33}$ (E), and APE1$^{K4pleA}$ (F).

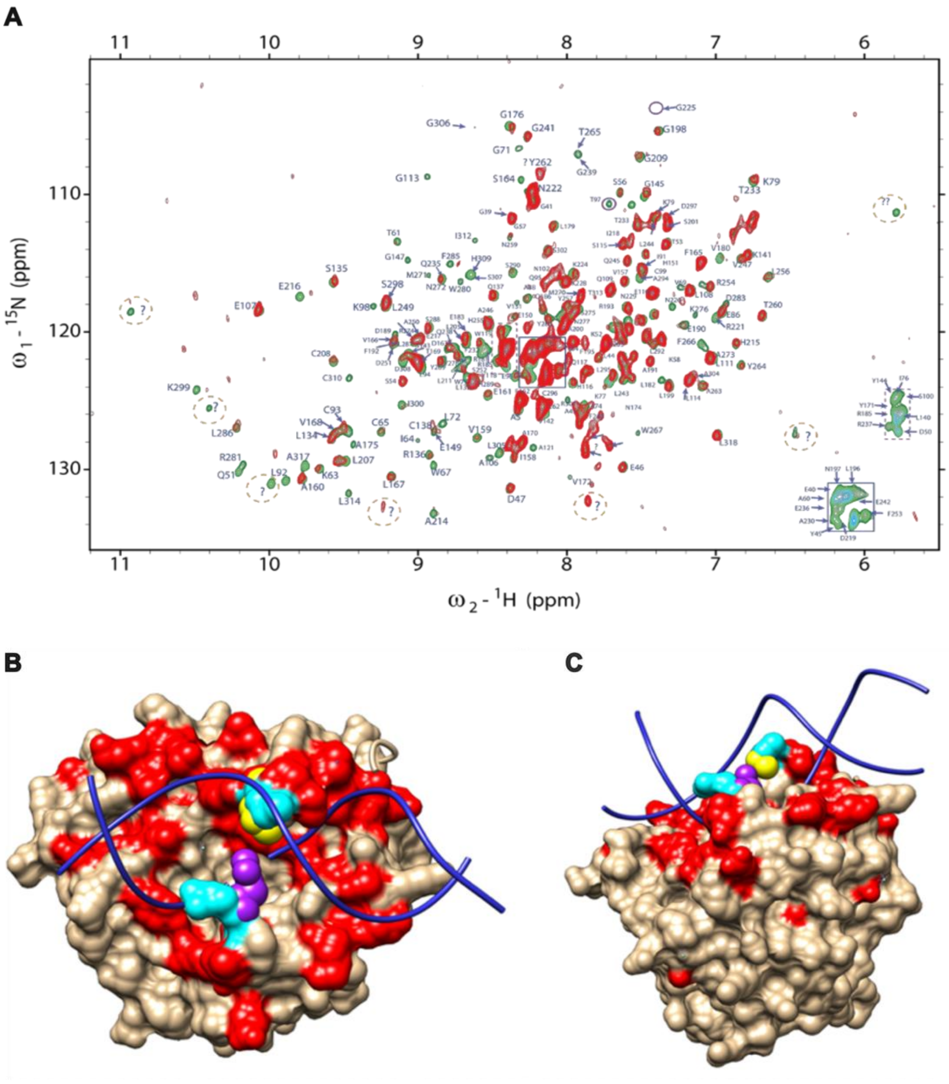


**Fig. S3.** A) 800 MHz $^{1}H$-$^{15}N$ TROSY-HSQC spectrum of $^{15}N$-labeled APE1 (green) and in presence of one molar equivalent of pre-92b oligoribonucleotide. Spectra collected at 25°C. Inset A and B were placed bottom right to facilitate overcrowded region visualization. Some cross-peaks intensities decrease due to binding with RNA molecule. In general, the attenuation is more important for the residues in the surface surrounding the canonical APE1 DNA binding site, close to R177. B-C) Visualization of the most important observed chemical shift changes, and as well the appearing and disappearing peaks due to different degrees of chemical exchange, indicating binding events. R177 and M270 make a bridge-like structure that inserts into the major and minor grooves of the DNA, respectively (purple for R177 and yellow for M270). The representation in red, of the most significative binding residues to pre-92b coincides with the canonical DNA binding site (blue ribbons).

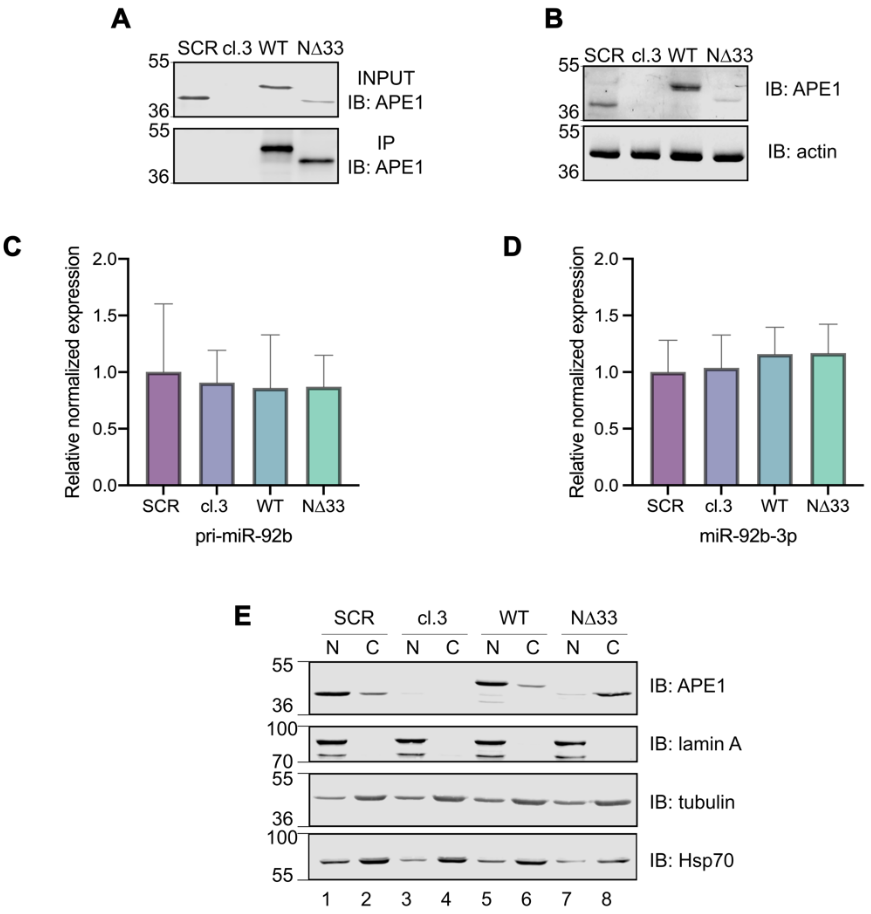


**Fig. S4.** A) Representative western blot analysis relative to RIP experiment showing APE1 on total HeLa cell clones extracts (INPUT) and on the immunoprecipitated material (IP). The different molecular weights are indicated in kDa on the left side of the panels. B) Western Blot analysis showing endogenous APE1 expression in HeLa clones. Staining with anti-actin antibody was carried out as a loading control. The different molecular weights are indicated in kDa on the left side of the panels. C) pri-miR-92b and D) miR-92b-3p levels in HeLa clones silenced for endogenous APE1 expression were evaluated by qRT-PCR. The histograms show the expression levels of pri-miR-92b, normalized to GAPDH levels and the expression levels of miR-92b-3p, normalized to miR-16-5p levels and normalized to APE1$^{WT}$. The data are expressed as means ± S.D. of three independent replicates. E) Representative WB analysis to confirm nuclei-cytosol fractions by loading nuclear and cytoplasmic HeLa clone extracts and incubated with anti-APE1, anti-lamin A, anti-tubulin, and anti-Hsp70 antibodies. The different molecular weights are indicated in kDa (MW).

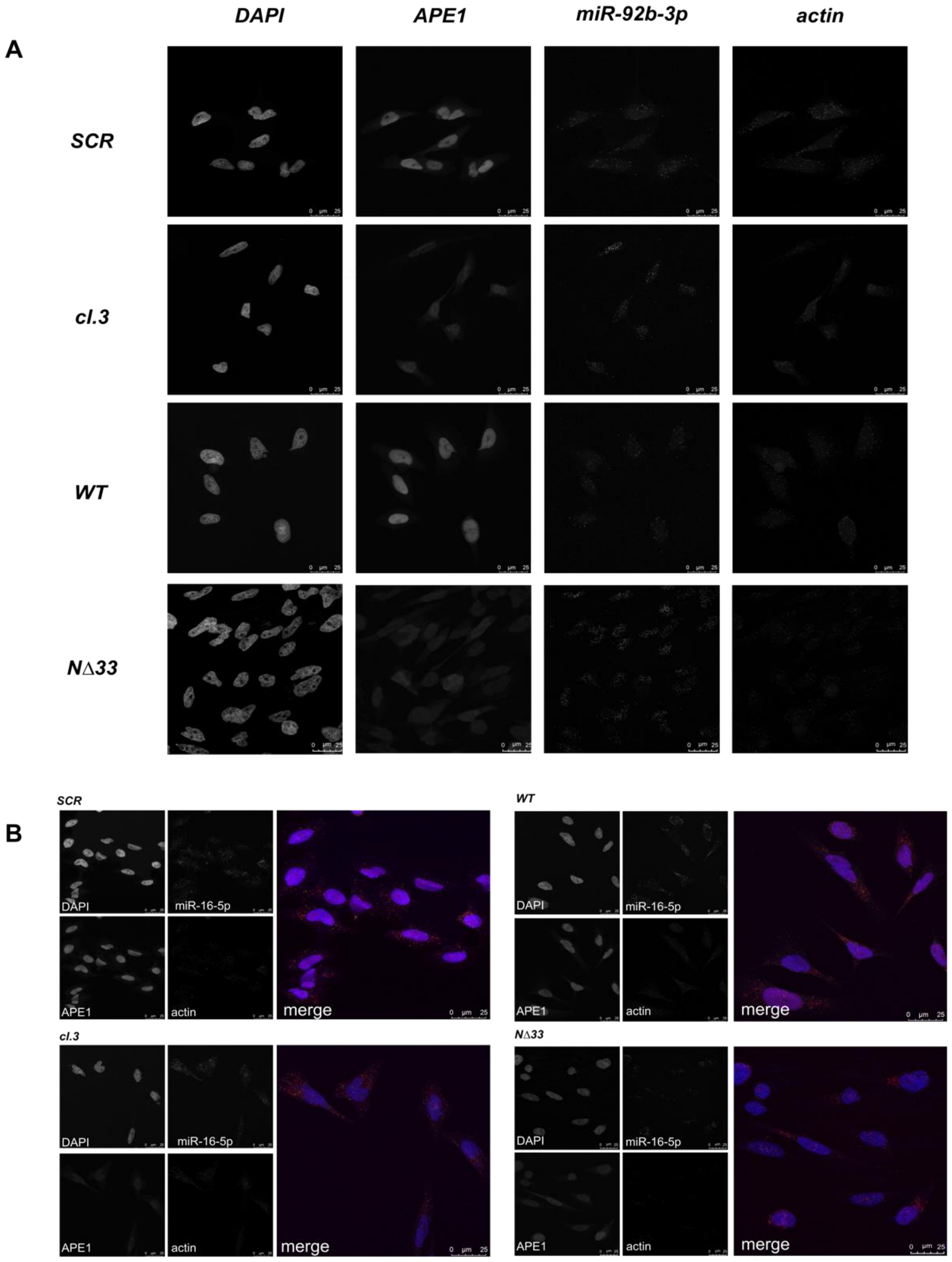


**Fig. S5.** A-B) Representative fluorescence confocal microscope images of FISH analysis for miR-92b-3p (A) and miR-16-5p (B) in HeLa clones. Cells were stained with probes specific for miR-92b-3p (A) and miR-16-5p (B) and actin mRNA, as control. APE1 staining was carried out using anti-APE1 specific antibody and nuclei were stained by DAPI.

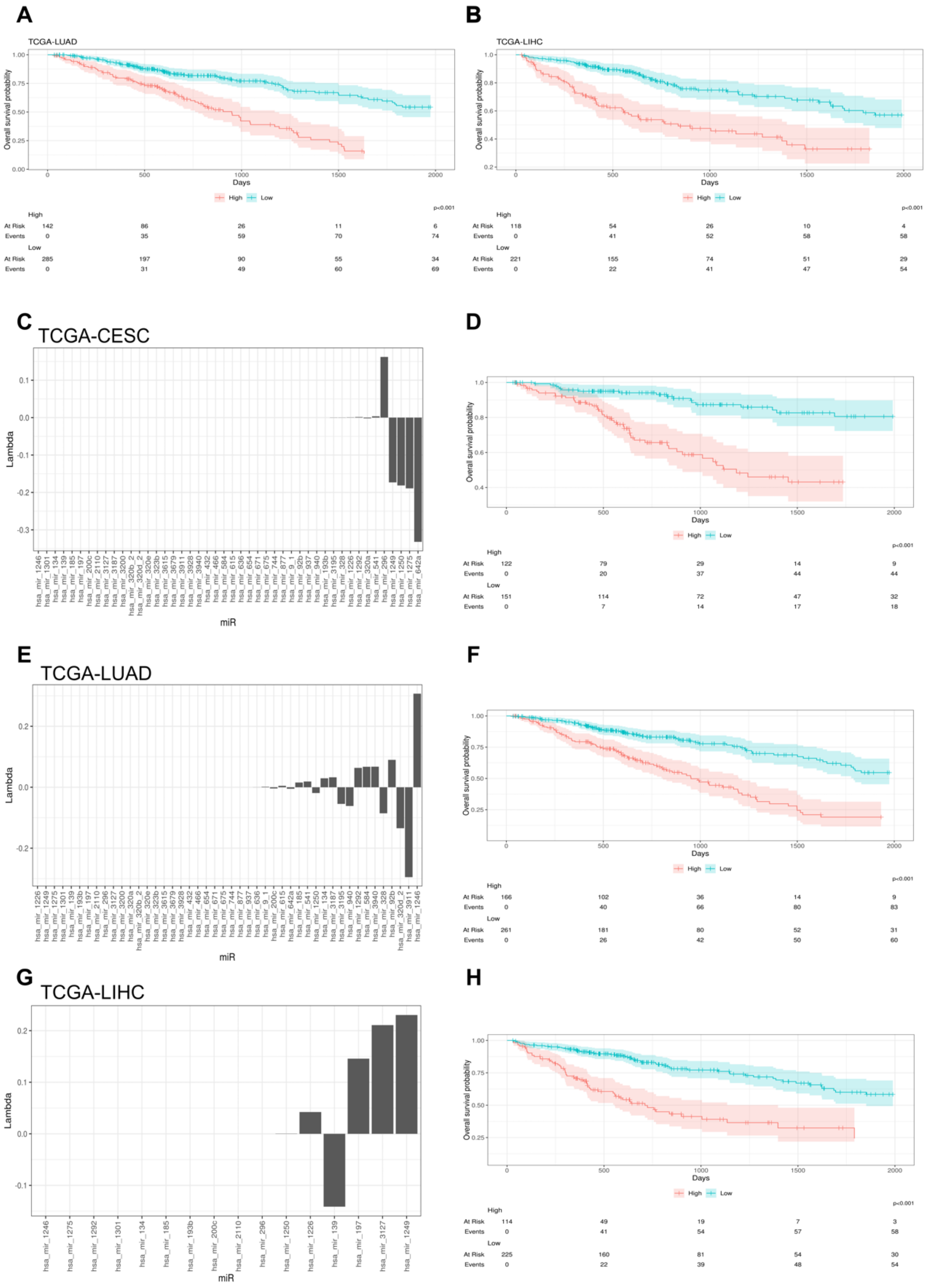


**Fig. S6.** Kaplan–Meier plots in TCGA-LUAD (A) and TCGA-LICH (B) dataset show the different OS rates of subjects belonging to the “High risk” and “Low risk” groups, stratified based on the PI calculated from the miRNA signature. Results have been reported from 0 to 2000 days as the sample size after 2000 days is too small to perform statistical inference. C-E-G) Bar plots reporting the lambda coefficients for each miRNA derived from the Cox model with LASSO penalization in TCGA-CESC (C), TCGA-LUAD (E) and TCGA-LIHC (G). miRNAs were ranked based on the absolute value of the lambda coefficients representing their associations with the OS rates. D-F-H) Prognostic value of the minimum miRNA signature in TCGA-CESC (D), TCGA-LUAD (F) and TCGA-LIHC (H) datasets. Kaplan–Meier plot showing the different OS rates of subjects belonging to the “High risk” and “Low risk” groups, stratified based on the PI calculated from the minimum miRNA signature. Associated *p-values* are reported under each plot. The number of samples in each dataset at different time points is reported in the table under the graph. Results have been reported from 0 to 2000 days as the sample size after 2000 days is too small to perform statistical inference.

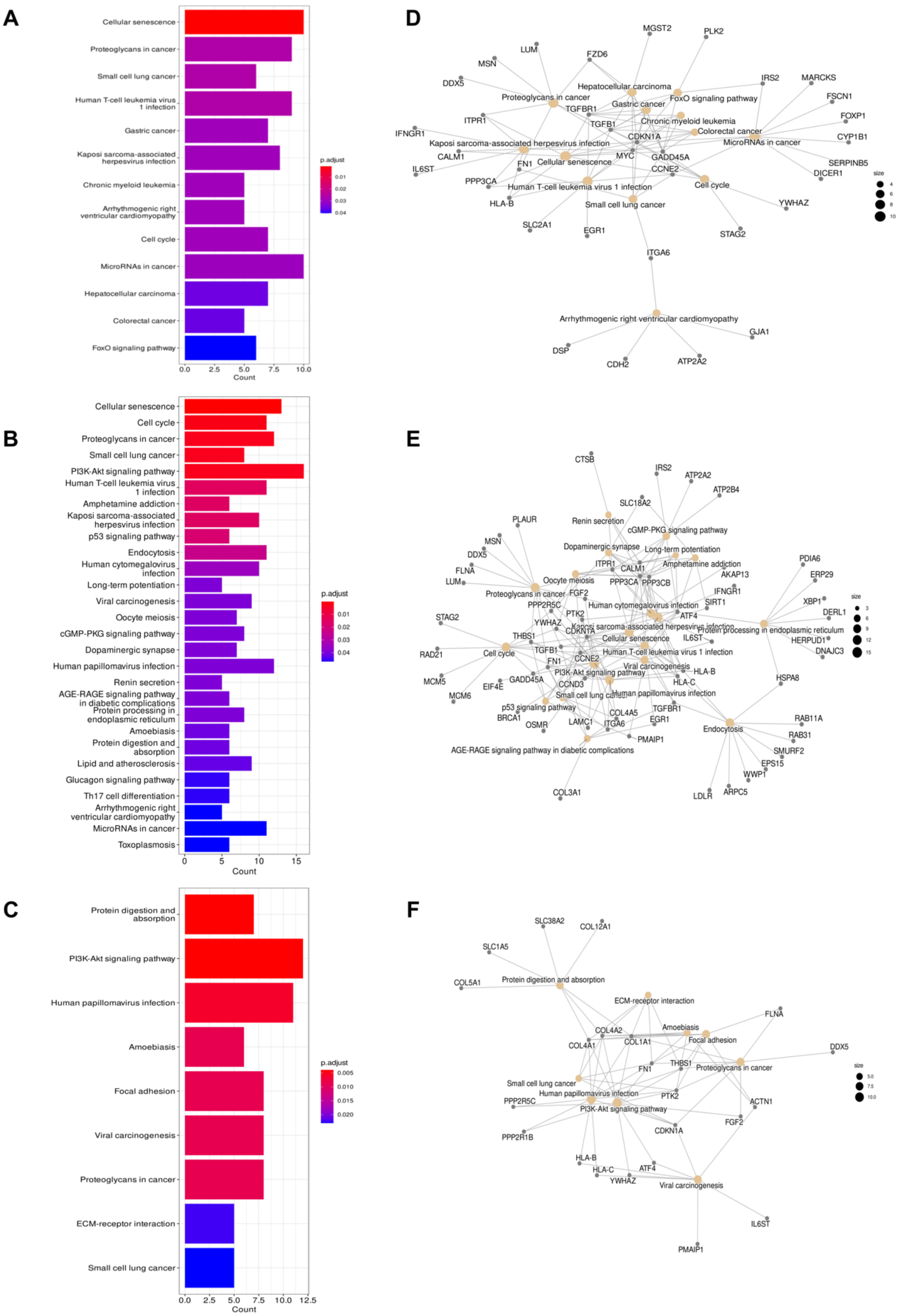


**Fig. S7.** A-C) Functional enrichment analysis of deregulated miRNA target genes showing the enriched KEGG pathways (adjusted *p-value* < 0.05) in the TCGA-CESC (A), TCGA-LUAD (B) and in the TCGA-LIHC (C) datasets. The colour of the bar represents the adjusted p-value while the x-axis the number of genes in each pathway. D-F) Representative network of the top 20 enriched KEGG pathways (adjusted *p-value* < 0.05) in the TCGA-CESC (D), TCGA-LUAD (E) and in the TCGA-LIHC (F) datasets.

**Table S1.** Potential rG4 forming sequences in pri-miR-92b (orange), pre-miR-92 (blue) and miR-92b-3p (purple), predicted with QGRSmapper tool. For each, the analyzed sequence and its length is reported. The QGRS output is also displayed, indicating the position of the G4 motif, the length, the sequence and the G-score.

**pri-miR-92b**

| **Analyzed sequence:** | **Length:** 196 |
|---|---|

GCCCCGGGUUCCCGGGUCGGGCCGGCUUCAGGCGGUGGGAGCGGGAUCCCGGCCC
CGGGCGGGCGGGAGGGACGGGACGCGGUGCAGUGUUGUUUUUCCCCCGCCAAUAUUG
CACUCGUCCCGGCCUCCGGCCCCCCCGGCCCCCCGGCCUCCCCGCUACCCCUAGCGGG
GCAGCCCCCGCCCUCCUGGCUCU

**QGRS output:**

| *Position* | *Length* | *Sequences (5′,3′)* | *G-Score* |
|---|---|---|---|
| *7* | *19* | ***GGUUCCCGGGUCGGGCCGG*** | *19* |
| *31* | *23* | ***GGCGGUGGGAGCGGGAUCCCGG*** | *21* |
| *59* | *15* | ***GGGCGGGCGGGAGGG*** | *42* |
| *126* | *26* | ***GGCCUCCGGCCCCCCCGGCCCCCCGG*** | *19* |

**pre-miR-92b**

| **Analyzed sequence:** | **Length:** 96 |
|---|---|

CGGGCCCCGGGCGGGCGGGAGGGACGGGACGCGGUGCAGUGUUGUUUUUCCCCCGC
CAAUAUUGCACUCGUCCCGGCCUCCGGCCCCCCCGGCCC

**QGRS output:**

| *Position* | *Length* | *Sequences (5′,3′)* | *G-Score* |
|---|---|---|---|
| *9* | *15* | ***GGGCGGGCGGGAGGG*** | *72* |

**miR-92b-3p**

| **Analyzed sequence:** | **Length:** 22 |
|---|---|

UAUUGCACUCGUCCCGGCCUCC

**QGRS output:**

| *Position* | *Length* | *Sequences (5′,3′)* | *G-Score* |
|---|---|---|---|
| ∅ | ∅ | ∅ | ∅ |

**Dataset S1 (separate excel file).** Table of dysregulated miRNAs after APE1 depletion in HeLa and A549 cellular models used for rG4 motifs prediction, obtained from Nanostring and RNA-seq experiments.

**Dataset S2 (separate excel file).** List of precursor and mature miRNAs holding an rG4 motif, predicted by QGRSmapper, pqsfinder, and G4hunter bioinformatic tools. miRNAs were listed considering their scoring, from the top hit to the bottom one. The threshold for the tools are respectively: QGRSmapper 19, pqsfinder 47, G4hunter 1.2.
